# Single Cell and Spatial Transcriptomics Defines the Cellular Architecture of the Antimicrobial Response Network in Human Leprosy Granulomas

**DOI:** 10.1101/2020.12.01.406819

**Authors:** Feiyang Ma, Travis K. Hughes, Rosane M.B. Teles, Priscila R. Andrade, Bruno J. de Andrade Silva, Olesya Plazyo, Lam C. Tsoi, Tran Do, Marc H Wadsworth, Aislyn Oulee, Maria Teresa Ochoa, Euzenir N. Sarno, M. Luisa Iruela-Arispe, Bryan Bryson, Alex K. Shalek, Barry R. Bloom, Johann E. Gudjonsson, Matteo Pellegrini, Robert L. Modlin

## Abstract

Granulomas are complex cellular structures comprised predominantly of macrophages and lymphocytes that function to contain and kill invading pathogens. Here, we investigated single cell phenotypes associated with antimicrobial responses in human leprosy granulomas by applying single cell and spatial sequencing to leprosy biopsy specimens. We focused on reversal reactions (RR), a dynamic process in which some patients with disseminated lepromatous leprosy (L-lep) transition towards self-limiting tuberculoid leprosy (T-lep), mounting effective antimicrobial responses. We identified a set of genes encoding proteins involved in antimicrobial responses that are differentially expressed in RR versus L-lep lesions, and regulated by IFN-γ and IL-1β. By integrating the spatial coordinates of the key cell types and antimicrobial gene expression in RR and T-lep lesions, we constructed a map revealing the organized architecture of granulomas depicting compositional and functional layers by which macrophages, T cells, keratinocytes and fibroblasts contribute to the antimicrobial response.

## Introduction

The hallmark of the chronic inflammatory response to a foreign substance that has resisted destruction by an acute inflammatory response is the granuloma. Granulomas have been defined as structures “which are formed by the immune-mediated recruitment of white blood cells, and particularly rich in macrophages”^1^. In the context of infectious diseases, the function of the granuloma is to sequester and degrade microbial pathogens that have evaded the early immune response.

Leprosy offers an attractive model to investigate the mechanisms by which the human immune system combats intracellular bacteria as the disease presents as a clinical/immunologic spectrum ^2^. The histologic characteristic of leprosy is the granuloma, containing macrophages that have been infected by the pathogen, *Mycobacterium leprae,* and lymphocytes. Because leprosy manifests as a spectrum of disease in skin, the dynamics are accessible to study, in contrast to tuberculosis granulomas. At one end of the disease spectrum, tuberculoid leprosy (T-lep) typifies the host’s antimicrobial response, which controls the pathogen: there are few lesions, *Mycobacterium leprae* bacilli are rare, and patients eliminate the infection. At the opposite end, lepromatous leprosy (L-lep) represents susceptibility to disseminated infection, with numerous skin lesions and abundant bacilli. This disease spectrum is dynamic, as patients may undergo a reversal reaction (RR), in which patients generally upgrade, either spontaneously or during chemotherapy, from the lepromatous to the tuberculoid pole.

The structure of granulomas is distinct across the spectrum of leprosy. The granulomas in T-lep contain a core of mature macrophages with occasional multinucleated giant cells. These granulomas are organized with a mantle zone at the periphery of the granuloma containing lymphocytes and characterized by fibrosis. Granulomas in RR lesions are histologically similar to those in T-lep with the presence of intercellular edema. In L-lep, the granulomas are disorganized, and immature lipid-laden macrophages are prominent with lymphocytes scattered throughout.

The study of leprosy lesions has provided insight regarding the host immune response to intracellular bacteria and the architecture of granulomas. Through various approaches, it has been possible to define functional subpopulations of human T cells ^3–6^ and macrophages ^7^, and their microanatomic distribution, as well as the patterns of cytokine secretion that influence the outcome of infections caused by pathogenic mycobacteria ^8–11^.

Given that the resolution of the granulomatous response requires destruction of the foreign invader, the antimicrobial mechanisms that result in the death of the pathogen are central to understanding how granulomas contribute to host defense. A few pathways have been identified by the study of human cells that can lead to an antimicrobial activity against intracellular mycobacteria. Through activation via TLRs and secretion of IFN-γ, the innate and adaptive immune systems trigger the vitamin D-dependent induction of the antimicrobial proteins encoded by *CAMP* and *DEFB4A* ^7,12,13^ T cells release antimicrobial proteins encoded by *GNLY* and *IL26,* which can enter infected macrophages and exert a direct antimicrobial activity ^4,5,14,15^. These human pathways are not present in mice, which utilize other mechanisms such as the release of nitric oxide to kill mycobacteria.

We and others have investigated antimicrobial pathways at the site of disease by immunohistology which identifies specific proteins but cannot differentiate whether the positive cell has secreted or taken up that protein. Microarrays and bulk RNA sequencing measure the transcript levels of genes encoding a protein at a global level, but do not identify the cell expressing that mRNA. The advent of single cell RNA-sequencing (scRNA-seq) provides an opportunity to associate transcripts with individual cells and construct cell-cell interaction networks ^16–19^. A further advance, spatial sequencing (spatial-seq), makes it possible to visualize the expression of mRNAs in the context of the tissue morphology. Here, we integrated scRNA-seq with spatial-seq to determine the structure of the organized granuloma in leprosy including the potential contribution of specific cell types to the antimicrobial response against an intracellular bacterium.

## Results

### Single cell RNA sequencing identifies 12 cell types in leprosy lesions

To study the cellular composition and cell-state differences between RR and L-lep, we performed single cell RNA-sequencing by Seq-Well ^20^ on skin biopsy specimens from five RR and five L-lep patients (**Supplementary Table 1**). We obtained 21,318 cells from 10 biopsy specimens, detecting an average of 741 genes and 3,556 transcripts per cell. To study the heterogeneity of these cells, we selected variable genes and performed UMAP dimensionality reduction and cell clustering using the R package Seurat ^21^. We then performed differential expression analysis to find the cluster markers and overlapped the cluster markers to canonical cell type defining signature genes. Ultimately, we recovered 12 primary cell types across all 10 samples (**Fig. 1a, b**). These annotated cell types include: T cells (TC; *CD3D* and *TRBC2*), B cells (BC; *MS4A1* and *CD79A*), plasma cells (PLC; *IGHG1* and *IGHG3*), myeloid cells (ML; *C1QA* and *LYZ*), Langerhans cells (LC; *CD207* and *CD1A*), mast cells (Mast; *CPA3* and *CTSG*), keratinocytes (KC; *KRT1* and *KRT10*), fibroblasts (FB; *COL1A1* and *DCN*), smooth muscle cells (SMC; *ACTA2* and *TAGLN*), endothelial cells (EC; *PECAM1* and *CDH5*), eccrine gland cells (ECG; *DCD* and *MUCL1*) and melanocytes (MLNC; *DCT* and *PMEL*) (**Fig. 1c, Supplementary Table 2**).

**Fig. 1.**
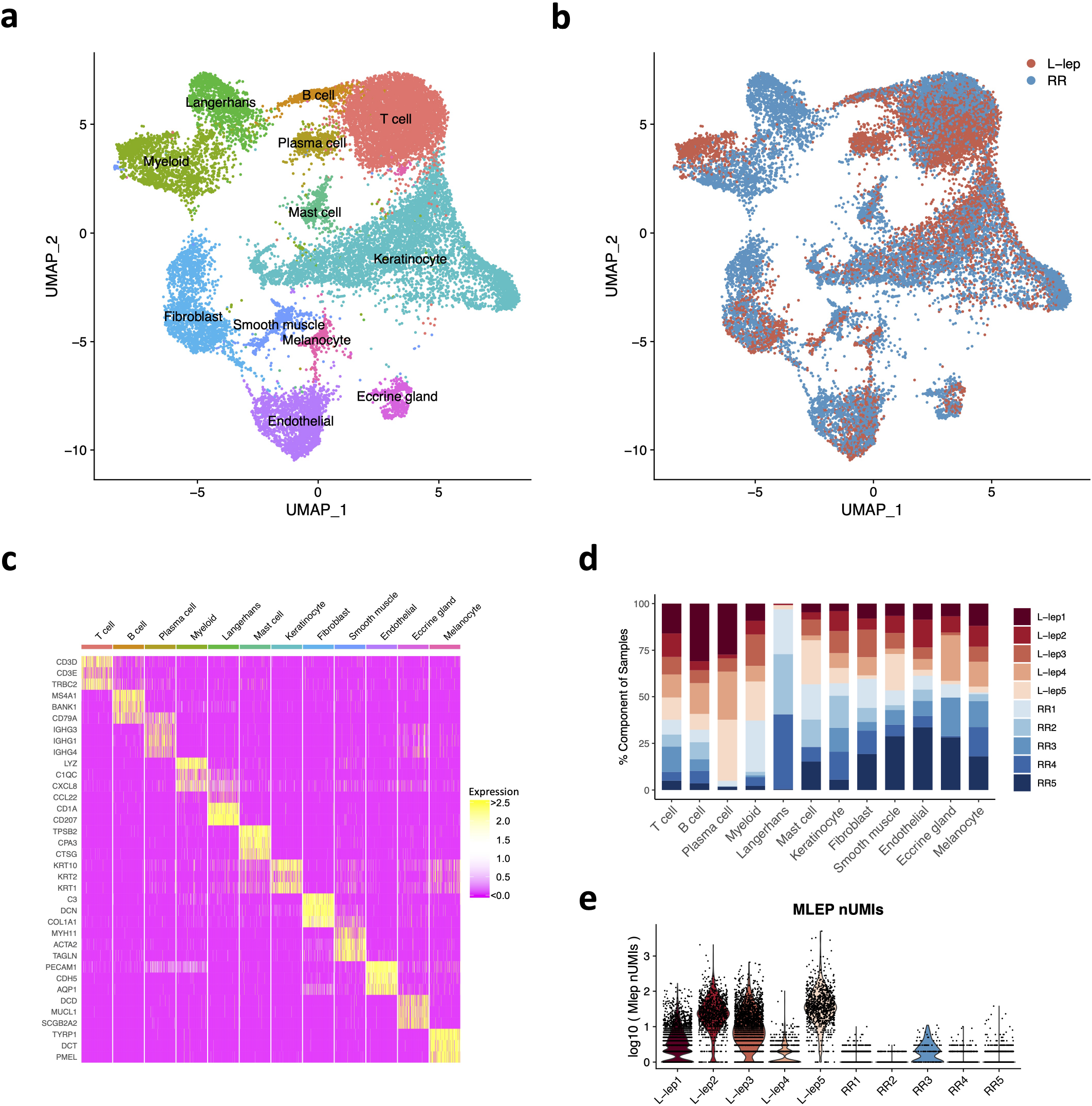
Cell types observed in leprosy lesions. a. UMAP plot for 21,318 cells colored by cell types. b. UMAP plot colored by clinical forms. c. Heatmap showing three representative marker genes for each cell type. d. Abundance composition across all samples for each cell type. e. Violin plot showing the number of *M. leprae* transcripts detected in individual cells from each patient.

The major cell types, including T cells, myeloid cells, keratinocytes, endothelial cells and fibroblasts, were found in both RR and L-lep lesions (**Fig. 1d**). Although B cells were found in both RR and L-lep lesions, plasma cells were derived predominantly from L-lep lesions. Given that LC are more frequent in RR than L-lep lesions ^3^, we isolated CD1a^+^ cells from the epidermis of three RR patients (RR1, RR2 and RR4) by immunoselection, adding these to the epidermal and dermal cells, accounting for the high frequency of LC from these RR lesions. *M. leprae* reads were most prevalent in the multibacillary L-lep lesions, but were also detected at a lower level in one RR lesion (**Fig. 1e, Supplementary Fig. 1**).

### Distinct cell subtypes in RR versus L-lep lesions differentiate the immune response in leprosy

We defined seven T cell sub-clusters, two predominantly derived from RR lesions and one from L-lep lesions, which we annotated using previously defined cluster specific genes (**Fig. 2a**) ^19^. T cell sub-cluster 0 (TC0), composed of >90% cells from RR samples, expresses the classic Th17 cell markers *RORC, RORA, RBPJ* and *IL23R* as signature genes (**Fig. 2b, c**), but not the major Th17 cytokine genes. TC1 and TC2 are designated as cytolytic T lymphocytes (CTL) as they both contain *CD8A, GZMB* and *PRF1.* Because TC1 was derived mainly from the RR lesions and TC2 was mainly derived from the L-lep lesions, they were labelled as RR CTL and L-lep CTL respectively (**Fig. 2d**). We noted several type I IFN downstream genes in L-lep CTL including *IFI44L, MX1, IRF1* and *OAS3.* We ran differential expression analysis between L-lep CTL and RR CTL, and performed enrichment analysis on the differentially expressed genes using IFN signatures derived from activated human PBMC ^11^. Genes up-regulated in L-lep CTL were significantly enriched in type I IFN downstream signatures (**Supplementary Fig. 2a, Supplementary Table 3**). The remaining sub-clusters contained a mixture of cells from RR and L-lep including TC3 (TCM, T-central memory; *IL7R* and *CCR7*), TC4 (naïve; *LEF1, JUNB*), TC5 (Treg; *FOXP3, CTLA4*) and TC6 (amCTL; antimicrobial CTL expressing *GZMB, PRF1* and *GNLY*) containing a mixture of tri-cytotoxic T cells (T-CTL) ^22^ and γδ T cells (**Fig. 2c**).

**Fig. 2.**
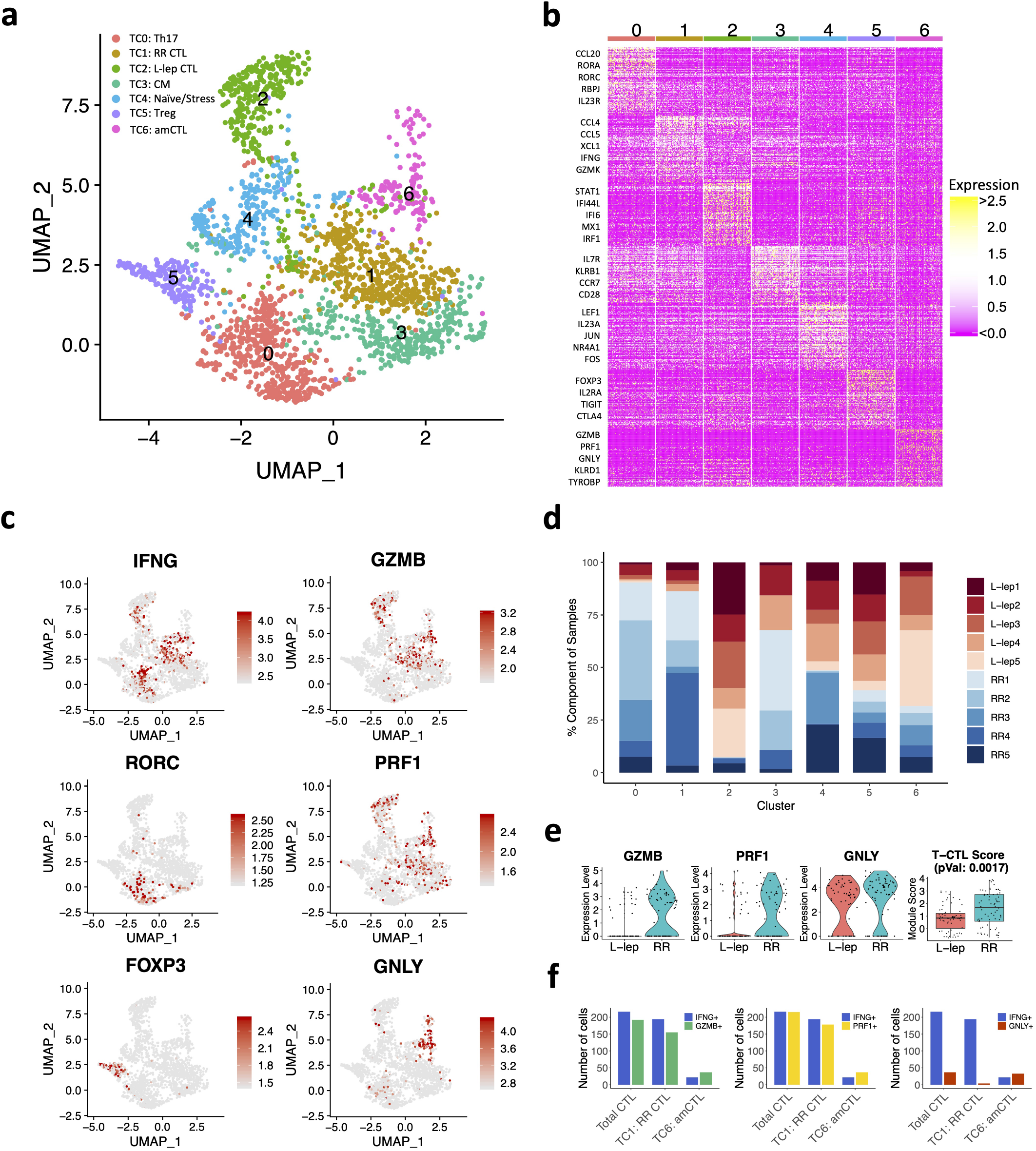
Identification of T cell subtypes. a. UMAP plot for 2,290 T cells colored by subtypes. b. Heatmap showing marker genes for each subtype. The representative genes are labelled. c. UMAP plots showing six marker genes. The color scale represents normalized expression level of the gene. d. Abundance composition across all samples for each T cell subtype. e. (Left) Violin plots showing the expression for *GZMB, PRF1* and *GNLY* in T cell sub-cluster 6 grouped by L-lep and RR. (Right) Boxplot showing the T-CTL score in T cell sub-cluster 6 grouped by L-lep and RR, the p value was calculated from a Wilcoxon rank sum test. f. Number of RR cells expressing *IFNG, GMZB, PRF1* and *GNLY* in RR CTL and amCTL.

We previously described a functional CTL subset, amCTL, expressing *GZMB, PRF1* and *GNLY,* that exert antimicrobial activity against intracellular *M. leprae* and correlate with protective immunity to tuberculosis and leprosy ^22,23^. In TC6, the expression of *GZMB, PRF1* and *GNLY* was greater in cells from RR vs. L-lep lesions. The module score aggregating the expression of these three antimicrobial CTL genes was significantly greater in RR lesions (**Fig. 2e**). *IFNG* was most strongly expressed by Th17 cells and RR CTL but was also present in amCTL and L-lep CTL (**Fig. 2c**). The frequency of *IFNG* expressing cells was similar to the frequency of *GZMB* and *PRF1* expressing cells in RR CTL and amCTL, but was far greater than the frequency of *GNLY* expressing cells in RR lesions (**Fig. 2f, Supplementary Fig. 2b**). These data indicate that *IFNG* is a marker for all CTL, but is not a useful marker for estimating amCTL, the small subset of CTL that express *GNLY,* in addition to *GZMB* and *PRF1,* and shown to kill the bacteria within infected cells.

We identified five myeloid sub-clusters, three predominantly derived from the RR lesions (ML0, ML3, ML4) and two from the L-lep lesions (ML1, ML2) (**Fig. 3**). ML0 (dendritic cells), containing cells from approximately 75% of RR lesions, is comprised of a mixture of dendritic cells with distinct subpopulations expressing *CD1C* and *LAMP3* (**Fig. 3a-d, Supplementary Fig. 3a**). ML1 (Type I IFN macrophages), containing >90% cells from L-lep lesions, are macrophages that express type I IFN downstream genes including *IFI44L, MX2* and *IFIT3,* hence categorized as Type I IFN macrophages (**Fig. 3c**). ML2, (TREM2 macrophages), containing >90% of cells from L-lep lesions, was characterized by high expression of *TREM2* and *APOE,* resembling the TREM2 macrophages reported in seven diseases characterized by altered lipid metabolism but not previously infectious disease ^24–30^. ML4, containing approximately 85% of cells from RR lesions, are M1-like macrophages, with *LYZ*, *MMP9* and *IL23A.* ML3 was derived from three RR lesions and two L-lep lesions, connecting TREM2 macrophages and M1-like macrophages (**Fig. 3b, d**), expressing genes from both sub-clusters (**Supplementary Fig. 3b**). *TREM2* and *APOE* expression, as well as a TREM2 score comprised of nine conserved genes from the datasets from the seven studies of lipid metabolism diseases, were highest in TREM2 macrophages, declining in transitional macrophages and M1-like macrophages (**Fig. 3e, Supplementary Fig. 3c**). These data indicate that TREM2 macrophages are predominantly found in L-lep lesions, with the expression of these signature genes declining with the transition to M1-like macrophages from RR lesions.

**Fig. 3.**
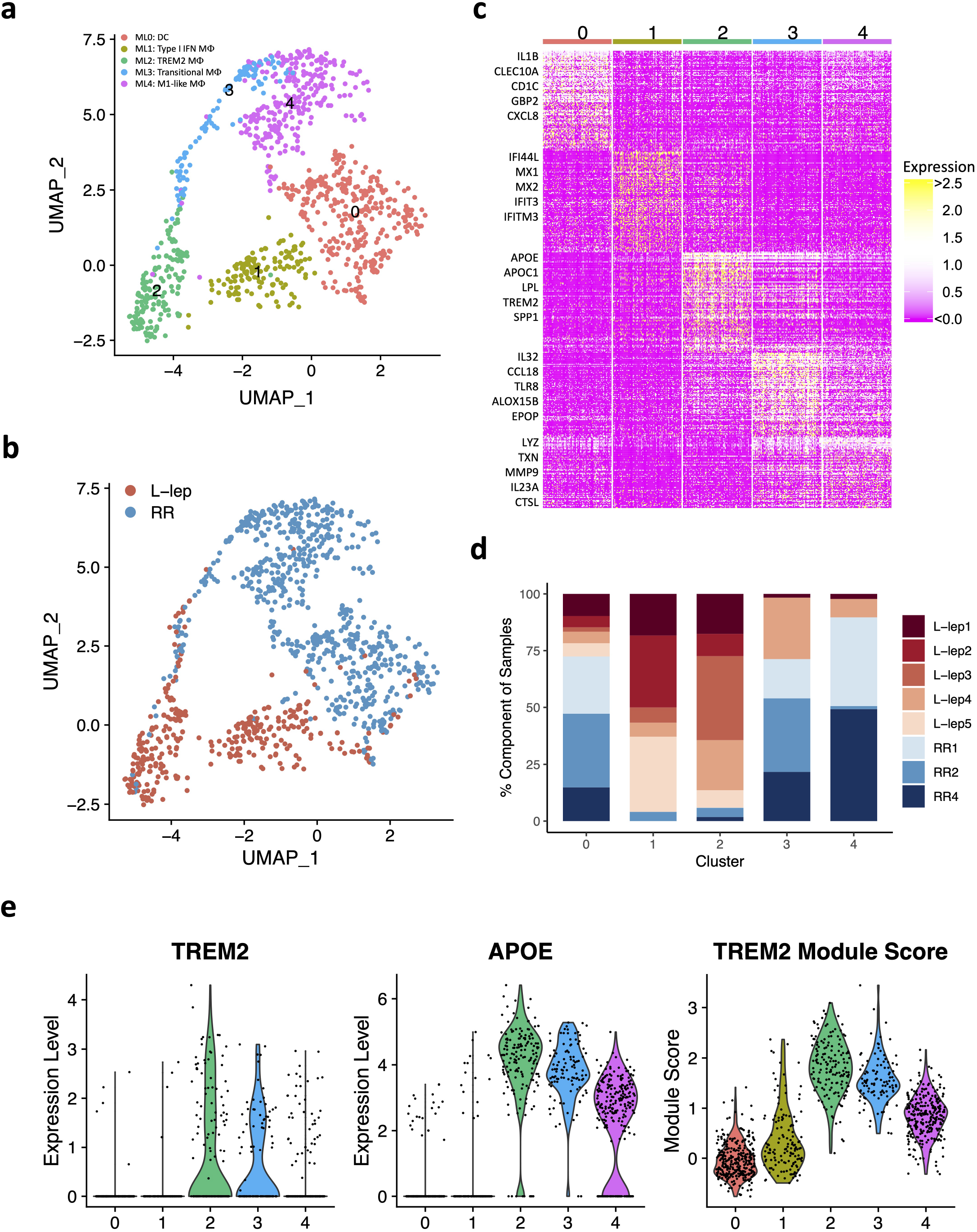
Identification of myeloid cell subtypes. a. UMAP plot for 991 myeloid cells colored by subtypes. b. UMAP plot colored by clinical forms. c. Heatmap showing marker genes for each subtype. The representative genes are labelled. d. Abundance composition across all samples for each myeloid cell subtype. e. (Left) Violin plots showing the expression for *TREM2* and *APOE* in myeloid subtypes. (Right) Violin plot showing the TREM2 Module score in myeloid subtypes.

Of seven keratinocyte sub-clusters, two were enriched in RR patients, KC3 (*FLG*^+^ granular layer keratinocytes) and KC4 (*KRT14/15* basal layer keratinocytes), and two enriched in L-lep patients, KC1 and KC5, both derived from spinous layer keratinocytes (**Supplementary Fig. 4a-d**). The RR enriched KC3 and KC4 clusters had increased expression of the proinflammatory gene *IL36G,* proteases *KLK5* (kallikrein 5) and *KLK7* (kallikrein 7), the protease inhibitor *PI3* (peptidase inhibitor 3 / elafin), the *CX3CL1* chemokine, and the antimicrobial *S100A2* (**Supplementary Fig. 4c**). By contrast, no increase was observed for either inflammatory or antimicrobial gene expression in the two L-lep keratinocyte clusters (KC1 and KC5). KC0 (spinous-1 keratinocytes), KC2 (supraspinous keratinocytes) and KC6 (hair follicle keratinocytes) were derived from both RR and L-lep samples.

*SFRP2^+^* fibroblasts (FB0) and *CXCL2^+^* fibroblasts (FB2) were enriched in RR lesions, with two additional sub-clusters mainly derived from L-lep lesions (**Supplementary Fig. 5**). The *SFRP2^+^* fibroblast sub-cluster is similar to a counterpart in normal skin, sharing expression of *COL3A1, COL18A1* and *COMP*^31^. *SFRP2^+^* fibroblasts from skin have been shown to have the capacity to release extracellular matrix proteins ^31^. The *SFRP2^+^* fibroblasts more strongly express genes that encode proteins involved in the deposition and modeling of the extracellular matrix including *VIM* (vimentin), *SPARC* (osteonectin), *FBN1* (fibrillin-1), *TNC* (tenascin), *FBLN2* (fibulin-2), *LOXL1* (lysyl oxidase homolog 1), as well as various collagens. By contrast, the *CXCL2*^+^ fibroblasts express a number of inflammatory genes as specific marker genes including *IL6*, *CCL2*, *CXCL3*, *CXCL8* and *IL32*, which displays a similar expression profile to the inflammatory fibroblasts detected in atopic dermatitis skin lesions ^32^. Two fibroblast sub-clusters were predominantly derived from L-lep lesions: *MGP^+^* fibroblasts (FB1) are a population of fibroblasts found in the reticular dermis and *COL11A1^+^* fibroblasts (FB3) have also been reported in skin ^31^.

Two endothelial cell sub-clusters, *LYVE1^+^* lymphatic EC (EC4) and *HEY2^+^* EC (EC5) were mainly derived from RR lesions. A previous study reported lymphatic malfunction in lepromatous leprosy ^33^, and our data suggests that lymphatic vessels were diminished in lepromatous lesions. *MEOX2*^+^ EC (EC3) were primarily found in L-lep lesions (**Supplementary Fig. 6**); *MEOX2* is an inhibitor of NF-κB activation in endothelial cells ^34^.

### Antimicrobial genes were highly expressed in RR cell types compared to L-lep cell types

Given that leprosy RRs are associated with a reduction in viable *M. leprae* bacilli in lesions, we sought to determine the array of antimicrobial genes that were present in defined cell populations. To do so, we first integrated a list of 1,404 “antimicrobial genes” curated by Genecards known to encode proteins that contribute to antimicrobial responses with the scRNA-seq data. We then separated each cell type (including subtype) into RR versus L-lep cells and calculated the sum of the expression of each gene across all cells of each cell type, and converted this sum to a z score (**Methods**). This metric captures the total amount of each transcript expressed by a cell type in our lesions, which measures the extent of the potential antimicrobial effect of its product. We compared the sum of z scores for L-lep and RR cell types and found that RR cell types have a higher expression pattern for the antimicrobial genes (**Fig. 4a**). We identified 1,124 antimicrobial genes with a z score ≥3 indicating robust levels of cell type specificity in at least one RR cell type. A high z score indicates that these cells, in aggregate, produce relatively more of the specific transcript than other cell types. We identified the upstream regulators of the 1,124 antimicrobial genes using Ingenuity Pathways Analysis (**Methods**), with *IFNG, TNF* and *IL1B* having the highest enrichment scores (**Fig. 4b**). The sum of the z scores for the top 20 upstream regulators of the antimicrobial genes was more highly expressed in the RR as compared to the L-lep cell types (**Fig. 4c**).

**Fig. 4.**
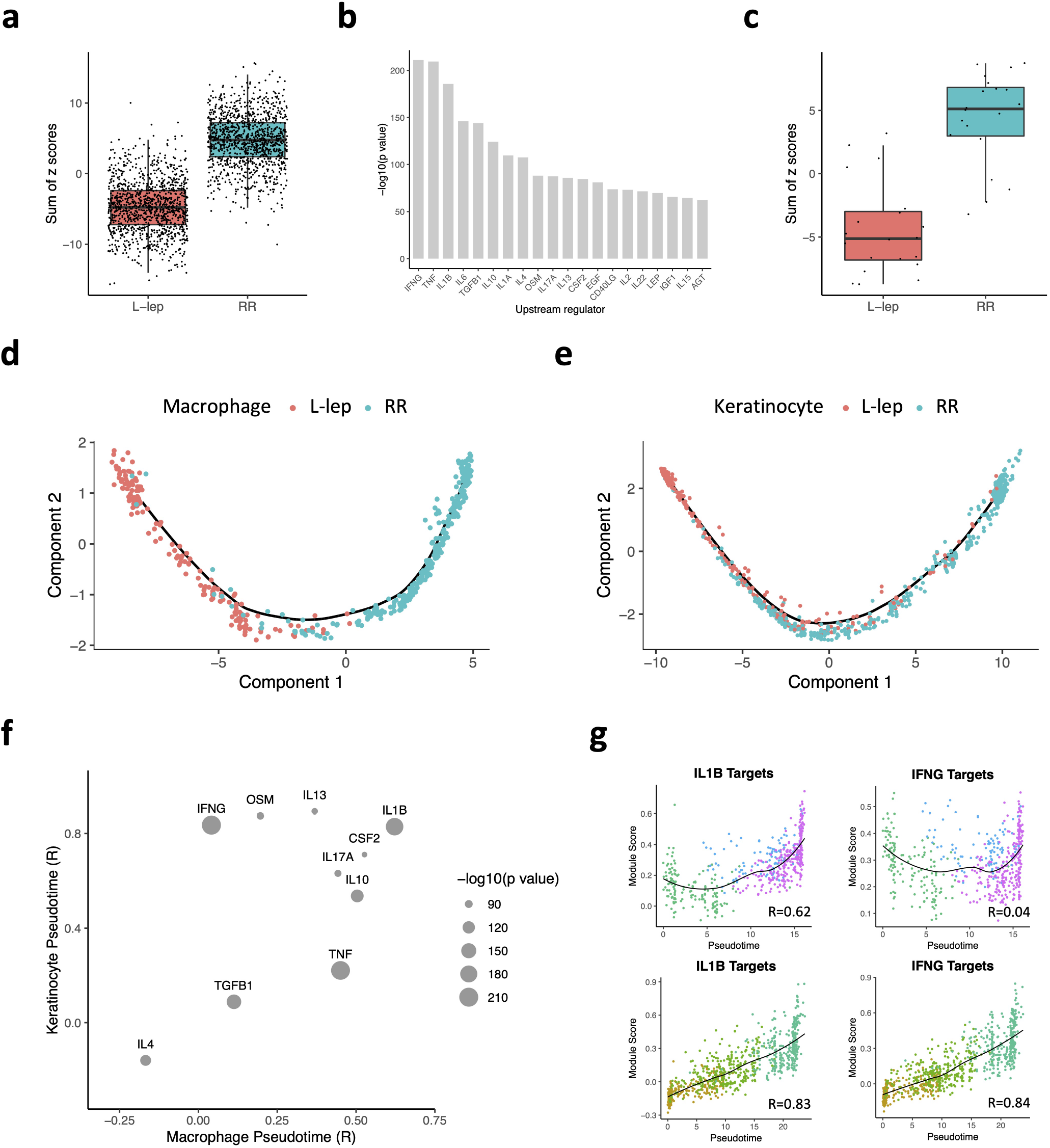
Antimicrobial gene analysis and pseudotime construction. a. Boxplot showing the sum of 1,124 antimicrobial gene z scores in L-lep and RR cell types. b. Bar graph showing the top 20 upstream regulators ranked by p value from the enrichment analysis using the 1,124 antimicrobial genes. c. Boxplot showing the sum of the z scores for the top 20 upstream regulators in L-lep and RR cell types. d. Pseudotime trajectory colored by clinical form in myeloid sub-cluster 2, 3 and 4. e. Pseudotime trajectory colored by clinical form in keratinocyte sub-cluster 1, 2 and 3. f. Dot plot showing the correlation between the module scores of the top 10 upstream regulators and macrophage/keratinocyte pseudotimes. The size of the dots represents the -log10(p value) from the enrichment analysis. g. Scatter plot showing the correlation between macrophage (top) or keratinocyte (bottom) pseudotimes and module scores calculated using *IL1B* target genes or *IFNG* target genes from the six expression patterns. Color of the dots represents the sub-cluster identity of the cells.

### Pseudotime analysis reveals macrophage and keratinocyte trajectories from L-lep to RR

Monocle, a pseudotime analysis program ^35^, created a trajectory in which cells were ordered from TREM2 macrophages (ML2), to ML3, terminating with M1-like macrophages (ML4) (**Supplementary Fig. 7a, b**). This trajectory starts from L-lep cells and ends with RR-derived cells, which mirrors the clinical progression seen in patients that start at the lepromatous pole and subsequently develop RR (**Fig. 4d**). Thus, ML3 appears to be transitional macrophages differentiating from TREM2 macrophage to M1-like macrophage. Using a similar analysis on spinous-2 KC (KC1), supraspinous KC (KC2) and granular KC (KC3), we identified a pseudotime continuum of the cells from these clusters (**Supplementary Fig. 7c, d**) that also progresses from L-lep to RR derived cells (**Fig. 4e**).

We hypothesized that the upstream regulators which trigger the antimicrobial response also drive the pseudotime trajectories in lesions. To test this, we split the variable genes into six expression patterns for both macrophage and keratinocyte pseudotimes (**Supplementary Fig. 7b, d, Supplementary Table 4**) and identified the upstream regulators for each expression pattern. We calculated a module score for each upstream regulator using the targets found in all six patterns and calculated the correlation coefficient between the module score and the pseudotime (**Methods**). Of the top upstream regulators for the antimicrobial genes, only the target scores for *IL1B* were highly correlated with both the macrophage and keratinocyte pseudotime (R=0.63 and 0.83, respectively) (**Fig. 4f, g**). The target scores for *IFNG* correlated with keratinocyte (R=0.84) but not macrophage pseudotime (R= 0.04). To this end, we selected for further study the *IFNG* and *IL1B* target genes, as *IFNG* had the highest enrichment score for the antimicrobial genes, and *IL1B* was not only a top upstream regulator of the antimicrobial genes but the expression of the *IL1B* target genes was highly correlated with both macrophage and keratinocyte pseudotimes.

### Cell networks reveal an antimicrobial ecosystem in RR lesions

To construct a gene network depicting the antimicrobial response in RR, we first determined the source of the two key upstream regulators. *IFNG* was detected (z score ≥3) in RR cells from Th17 cells and RR CTL, and *IL1B* was detected in RR cells from Langerhans cells and dendritic cells (**Fig. 5a**). As such, our analysis reveals that both the innate and adaptive immune systems contribute to the antimicrobial response by expression of *IL1B* and *IFNG*, respectively.

**Fig. 5.**
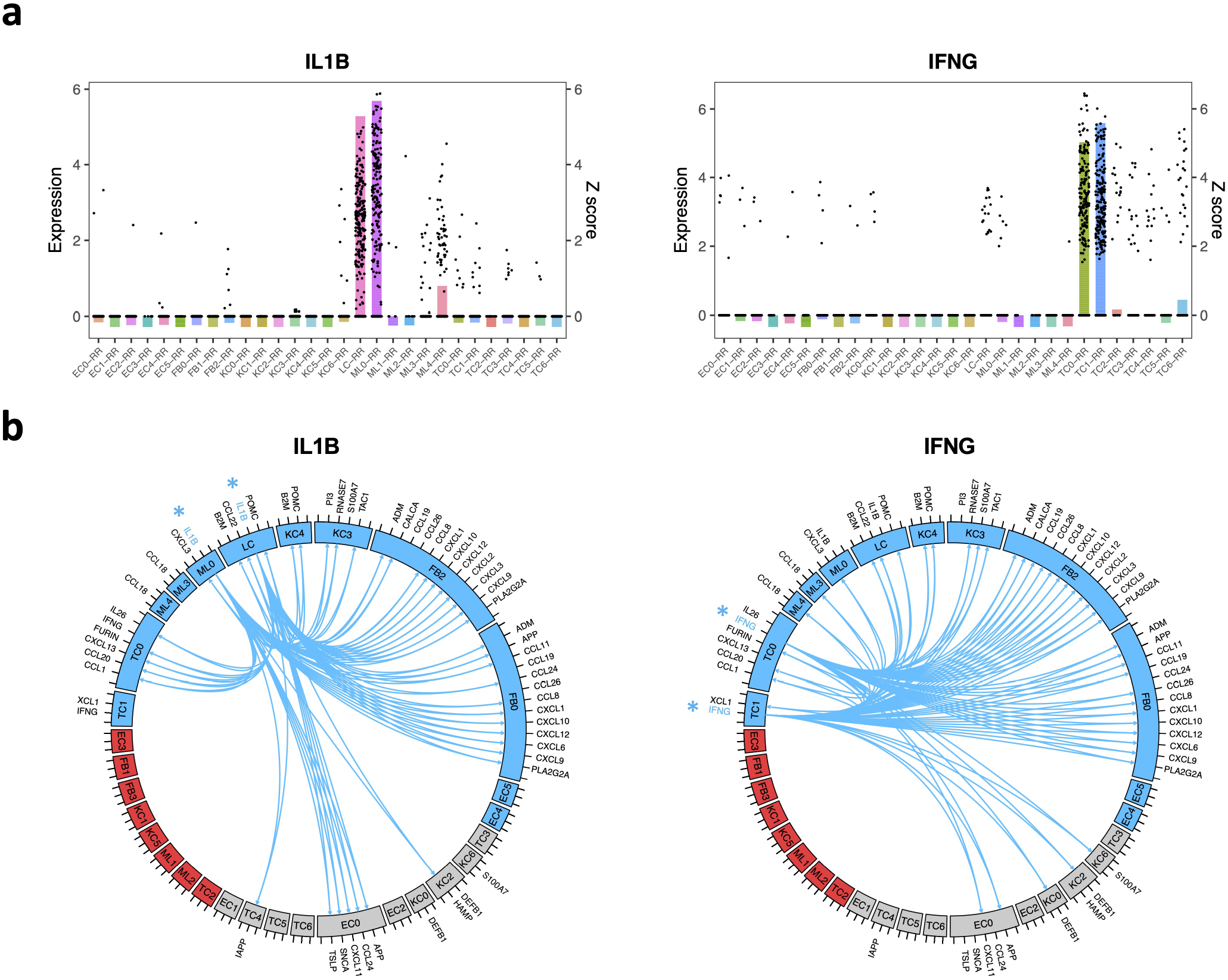
Antimicrobial network induced by *IL1B* and *IFNG.* a. Bar plot showing the z scores of *IL1B* (left) or *IFNG* (right) expression levels in each cell type from RR lesions. The dots represent *IL1B* or *IFNG* expression level in individual cells. b. Circos plot showing the connection between *IL1B* (left) or *IFNG* (right) and the direct antimicrobial gene targets in the cell types with z score >3. The color represents patient composition of the cell type. Red: L-lep specific; Blue: RR specific; Grey: mix of L-lep and RR.

Next, we constructed circos plots to depict the interactions of the *IL1B* and *IFNG* expressing cells with the target antimicrobial gene expressing cells (**Fig. 5b**). For clarity, we limited the number of interactions to the 61 antimicrobial genes that encode proteins with direct antimicrobial activity and with a z score ≥3 in at least one of the major RR cell types (endothelial cells, fibroblasts, keratinocytes, myeloid cells and T cells; **Supplementary Fig. 8 and 9**). In view of the variable expression of the genes encoding the receptors for IL-1β and IFN-γ, we inferred connections between the upstream regulators and these antimicrobial genes as identified using Ingenuity Pathway Analysis. *IL1B* was linked to 30 unique direct antimicrobial gene targets with 42 connections to RR cells, and *IFNG* was linked to 28 unique direct antimicrobial gene targets with 44 connections to RR cells. *IL1B* and *IFNG* both regulated 22 antimicrobial gene targets and individually regulated 14 additional antimicrobial genes, for a total of 36 unique genes. For both *IL1B* and *IFNG,* the majority of connections were to cell types that were predominantly derived from RR lesions, and strikingly there were no connections to a cell type that was predominantly derived from L-lep lesions. Although we had determined that *CXCL2^+^* fibroblasts expressed several chemokine genes that were cluster markers, the circos plot indicated that the *SFRP2^+^* fibroblasts also expressed several chemokine genes that encode for proteins with antimicrobial activity.

In an effort to validate the relevance of antimicrobial genes upregulated in RR lesions, we identified 59 genes that have been reported to be contribute to host defense against leprosy and other mycobacteria (**Supplementary Table 5, S6**). Of the 59, 34 genes have been reported to encode for proteins with direct antimicrobial activity ^36^. There were an additional seven genes reported as directly antimicrobial that had not previously been reported to be involved in the immune response to mycobacteria. We plotted the z scores for these 66 antimicrobial genes in a heat map arranged according to the specific cell type in which the antimicrobial gene was most strongly expressed (**Supplementary Fig. 10**). Strikingly, we found that each major cell type highly expressed a distinct set of antimicrobial genes. For example, *CYBB* was predominantly expressed in M1-like macrophages (ML4) and LCs, *CCL5* was predominantly expressed by RR CTL (TC1), *MMP2* was predominantly expressed by *SFRP2*^+^ fibroblasts (FB0) and *CXCL2^+^* fibroblasts (FB2), and *KLK5* was predominantly expressed by granular keratinocytes (KC3). Less frequently, antimicrobial genes such as *CXCL10* were expressed by multiple cell types.

### Spatial sequencing identifies the coordinates of cell populations in leprosy lesions

To localize the various cell types and antimicrobial responses within the granuloma architecture, we performed spatial-seq on leprosy skin biopsy specimens. We were able to collect RNA from a 20μm thick frozen section of an RR patient into 55μm wells containing spatially-barcoded capture oligonucleotides and performed transcriptome analysis. We detected 708 spatially-defined spots with an average of 2,984 genes and 10,007 transcripts per spot. The RR biopsy specimen contained approximately 15 discrete granulomas throughout the dermis (**Fig. 6a**).

**Fig. 6.**
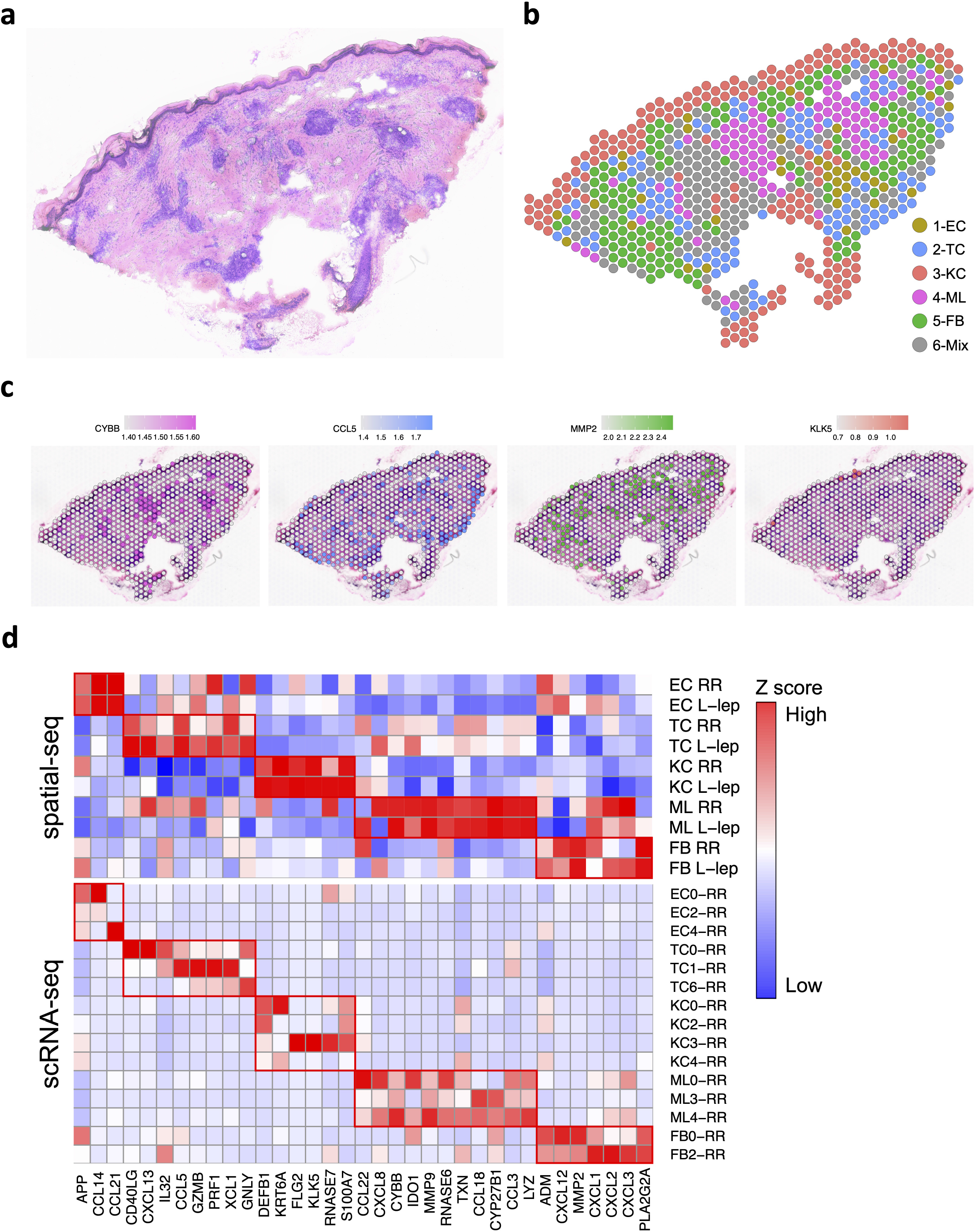
Spatial sequencing recapitulates the same set of antimicrobial genes in the corresponding cell types. a. H & E staining of the RR biopsy used for spatial sequencing. b. Spatial plot for 708 spots colored by clusters, the coordinates of the spot correspond to the location in the tissue. c. Spatial plots showing expression level of four antimicrobial genes. The colors correspond to the cluster colors in B. *CYBB* was highly expressed in myeloid cells, *CCL5* in T cells, *MMP2* in fibroblasts and *KLK5* in keratinocytes. d. Heatmaps showing 35 antimicrobial genes expressed in the same cell types in both spatial-seq and scRNA-seq. Both the RR and T-lep spatial-seq samples were plotted. The red boxes indicate distinct sets of antimicrobial genes highly expressed in a certain cell-type.

We annotated the cell-type composition for each spot by decomposing the spatial gene expression using the scRNA-seq gene expression of the major cell-types, displayed as a scatter-pie plot, which contains a pie chart for each spot scattered across the spatial array (**Supplementary Fig. 11a**). Thus, we identified the distribution of the five major cell type clusters revealing distinct aggregates of myeloid cells, T cells and fibroblasts in the dermis (**Fig. 6b, Supplementary Table 7**), and calculated the cell type composition scores for each cluster (**Supplementary Fig. 11b**). As RRs represent a transition from the lepromatous to the tuberculoid pole, we also performed spatial-seq on a skin biopsy specimen from a patient with T-lep. By histologic examination, this biopsy specimen contained more than ten granulomas (**Supplementary Fig. 12a**). The cell-type composition predicted from the scRNA-seq data in the T-lep lesion was similar to the RR lesions (**Supplementary Fig. 12b-d**).

We also sought to localize the RR cell subtypes in the morphologic context of the granuloma within the leprosy skin lesion. Strikingly, in both the RR and T-lep biopsy specimen, *SFRP2^+^* fibroblasts (FB0) were located in the superficial dermis near the dermal-epidermal junction, whereas *CXCL2*^+^ fibroblasts (FB2) were located deeper in the dermis around the granulomas (**Supplementary Fig. 13a, b**). The spatial-seq data validates the predictions of the epidermal layers from the scRNA-seq data for granular keratinocytes (KC3), spinous keratinocytes (KC0) and basal keratinocytes (KC4). The transitional macrophages (ML3) and M1-like macrophages (ML4) localized to the two large myeloid aggregates with dendritic cells (ML0) at the periphery.

Spatial mapping of four representative antimicrobial genes with defined distributions among the major cell-type sub-clusters in the scRNA-seq data (**Fig. 6c**), confirmed that the spatial localization of each gene mapped to the location of the corresponding cell type in the RR and T-lep spatial-seq data. *CYBB* expression was strongest in the same two aggregates of spots that were identified as myeloid cells, including transitional macrophages (ML3) and M1-like macrophages (ML4). *CCL5* was localized to regions that were defined by spatial-seq as containing T cells, with a mixture of T cell subpopulations. *MMP2* was detected in the spatial regions that contain fibroblasts, both FB0 and FB2. Finally, *KLK5* was expressed in the keratinocytes in the epidermis, in the region corresponding to granular keratinocytes (KC3), particularly in the T-lep sample.

We also investigated the expression of the 66 genes encoding proteins contributing to the antimicrobial response identified in the scRNA-seq data (**Supplementary Fig. 10**), finding that 55 of these genes were detected by spatial-seq of the RR and T-lep lesions. Of the 55 antimicrobial genes, 35 had high expression in the corresponding cell types in both the scRNA-seq and the spatial-seq data (**Fig. 6d**, **Supplementary Table 6**), with the remaining 20 genes having low or moderate expression in the corresponding spatial-seq cell type. There were 11 genes that were highly expressed in myeloid cells in both the scRNA-seq and spatial-seq data including *CYBB, CCL3, CCL18* and *MMP9.* Thus, the spatial-seq data validated the antimicrobial gene expression identified in the scRNA-seq data.

### Spatial sequencing maps the architecture of granulomas

The T-lep lesion contained a classic organized granuloma in the dermis of the biopsy specimen, composed of a core of macrophages surrounded by a mantle of aggregated lymphocytes (**Fig. 7a**). The spatial-seq data validated the histology for this region, showing the myeloid aggregates surrounded by T cells as well as regions of fibroblasts in the superficial dermis and around the myeloid aggregates. Close examination of the scatter-pie representation of this region identified some heterogeneity in the composition of the spots. For example, some spots in the epidermis were predicted to contain keratinocytes and a small proportion of Langerhans cells. Although the myeloid cells, T cells and fibroblast aggregates contained spots that were predominantly represented by their respective cell type, they were also composed of cells from the other cell types.

**Fig. 7.**
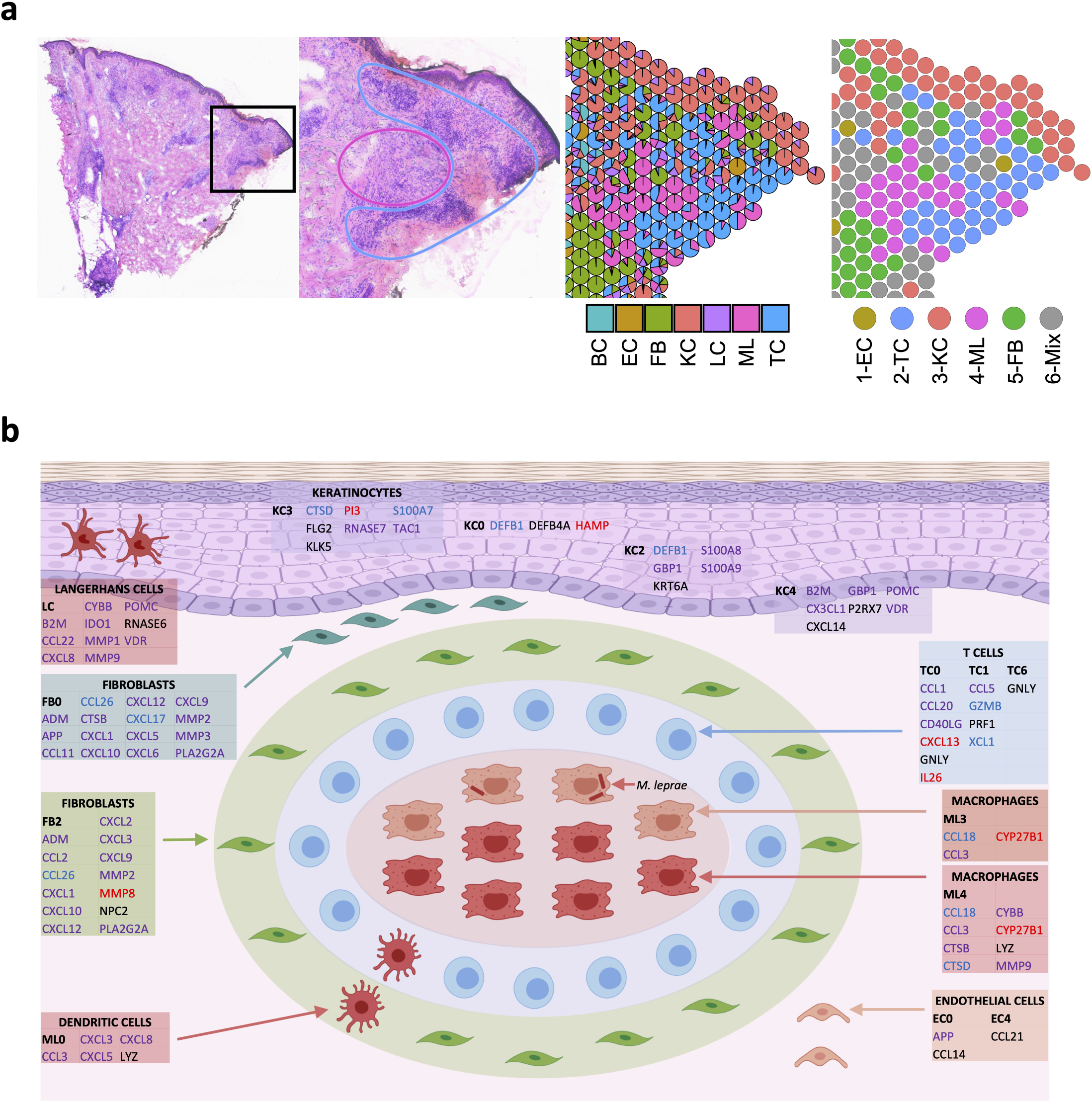
Granuloma architecture and antimicrobial ecosystem obtained by scRNA-seq and spatial-seq. a. H & E staining of the T-lep biopsy and the cell type composition highlighting the myeloid cells in the center of the granuloma and the T cells and fibroblasts at the periphery. b. Granuloma architecture and antimicrobial ecosystem. Gene names in red represent targets of *IL1B.* Gene names in blue represent targets of *IFNG.* Gene names in purple represents targets of both *IL1B* and *IFNG.* Gene names in black were curated from known antimicrobial pathways.

Thus, by integrating the scRNA-seq and spatial-seq data, we were able to create a model of the organized granuloma structure defined by cell subtypes and antimicrobial pathways (**Fig. 7b**). The clinical RR syndrome that develops in lepromatous patients is characterized by a change from immature to mature macrophages with a reduction in the number of intracellular bacilli. The model of the organized granuloma is composed of a core of macrophages, both transitional macrophages (ML3) and M1-like macrophages (ML4) that expressed genes encoding proteins with directly antimicrobial activity (*CCL3*, *CCL18*), as well as enzymes that contribute to the synthesis of antimicrobial mediators including *CYBB* and *CYP27B1* which facilitates the vitamin D-dependent induction of *CAMP* and *DEFB4A.* The adaptive T cell response was localized predominantly in the mantle zone, contributing to the antimicrobial response against intracellular bacteria in macrophage via amCTL expressing *GZMB*, *PRF1* and *GNLY*, RR CTL expressing *CCL5* and *XCL1* as well as Th17 cells expressing *IL26.* Also, surrounding the granuloma are dendritic cells (ML0) and *CXCL2^+^* fibroblasts (FB2), which expressed genes that encoded proteins with direct antimicrobial activity. Throughout the specimen were *SFRP2*^+^ fibroblasts (FB0) in the superficial dermis, various keratinocyte subtypes and endothelial cells, all expressing genes encoding antimicrobial proteins as well as chemokines known to have a direct antimicrobial response. The antimicrobial response in the granuloma and the surrounding microenvironment was largely regulated by *IL1B* and *IFNG.* Together, this suggests that the granuloma is a structured cellular ecosystem comprised of defined strata of distinct cell types that have the capacity to act to contain infection by an intracellular bacterium through the engagement of cells, both within and outside the granuloma, resulting in an integrated antimicrobial network.

## Discussion

The organized granulomatous response allows the immune system to wall off and eliminate intracellular bacteria that have initially evaded destruction. Across granulomas of leprosy and tuberculosis there is a spectrum, from spatially organized granulomas which can sterilize or contain the pathogen, as in T-lep and healed tuberculosis lesions, to those in which the pathogen flourishes and produces tissue damage, as in L-lep and miliary or progressive tuberculosis. Investigation of the immune interactions in such granulomas has over the past several decades, and almost exclusively, focused on the role of specific myeloid and lymphocytic populations. Our premise has been that the study of the changes in the cellular response in disease lesions of immunologically unresponsive patients with L-lep to the immunologically reactive patients with RR will enable understanding of mechanisms by which granulomas contribute to the antimicrobial response. By scRNA-seq, we identified 66 key antimicrobial genes that had been previously reported to contribute to host defense in leprosy and other mycobacterial infections, and/or encode proteins with direct antimicrobial activity. By integrating scRNA-seq with spatial-seq, it was possible for the first time to delineate the structure of the organized granuloma in leprosy that contributes to protective immunity according to the identify and locations of cell subtype and their antimicrobial capability.

At the core of the organized granuloma in the RR and T-lep lesions, is an aggregate of mature macrophages known as epithelioid cells, which we found were a mixture of M1-like macrophages (ML4) and transitional macrophages (ML3). The pseudotime trajectory analysis of the leprosy scRNA-seq data suggests the differentiation of TREM2 macrophages (ML2), predominantly found in L-lep lesions, to transitional macrophages (ML3), found in both RR and L-lep lesions, terminating in M1-like macrophages (ML4), found predominantly in RR lesions. We identified *IL1B* and *IFNG* as key upstream regulators of this pseudotime trajectory as well activation of macrophages in granulomas to express genes with the capacity to contribute to the antimicrobial response including *CCL3, CCL18, CYBB* and *MMP9,* located in the macrophage core of the granuloma in RR and T-lep lesions. Thus, the core of the protective granuloma contains differentiating and/or activated macrophages armed with multiple antimicrobial weapons in the battle against intracellular bacteria. The detection of an *IL1B* and *IFNG* triggered macrophage differentiation and activation program in leprosy lesions is consistent with the dynamic change from M2-like to M1-like macrophages during RR ^7^. Previously, we found that activation of TLR2, which like IL-1β signals via MyD88, in combination with IFN-γ, triggered the repolarization of M2-like into M1-like macrophages in vitro ^7,37^

The organized granuloma in T-lep patients was noted by Ridley and Jopling in their classification of leprosy to exhibit a peripheral zone of lymphocytes around the macrophage core ^2^, which was visualized in the spatial-seq analysis of RR and T-lep lesions. This mantle zone also contained dendritic cells (ML0) and as previously shown Langerhans cells ^3^, thus providing signals for T cell activation and differentiation, as well as *IL1B* which we identified as a key regulator of the antimicrobial response. We identified that various T cell subtypes were present in the mantle zone, including RR CTL and amCTL, which expressed *IFNG,* also a key regulator of the antimicrobial gene response, consistent with previous findings demonstrating that there is a dynamic change from a Th2 to a Th1 response including IFN-γ in paired samples in the same individuals from before and during RR ^8,10,11,33^ Therefore, the mantle zone of the granuloma has the capacity to secrete cytokines that trigger a downstream antimicrobial response in macrophages located in the core of the granuloma, as well as other distally located cells.

The mantle zone of T cells surrounding the macrophage core of an organized granuloma additionally contributes to host defense by the release of antimicrobial proteins that serve to contain bacteria trying to escape the granuloma following release from killed infected macrophages. The various T cell subpopulations detected in the mantle zone also express genes encoding proteins with direct antimicrobial activity. Th17 cells in RR lesions express *IL26,* which encodes an antimicrobial protein that has direct activity against intracellular *M. leprae* ^15,39^. Although amCTL were present in both RR and L-lep, there was higher expression of *GZMB, PRF1* and *GNLY* in the RR cells; these genes encode proteins that act synergistically when released from amCTL to kill intracellular mycobacteria ^4–6,22,23^. amCTL included γδ T cells, which we previously found were more frequent in RR lesions ^40^ and have been shown to release granulysin, thus able to kill infected macrophages and the mycobacteria within them ^41^. Among the two CTL sub-clusters, RR CTL and amCTL, the RR cells in both contained *IFNG* expressing cells, but only amCTL expressed *GNLY.* RR CTL were more abundant than amCTL, such that detection of IFN-γ is an appropriate surrogate measure of all CTL, but not of the specialized amCTL, which require expression of *GZMB, PRF1* and *GNLY* for their antimicrobial activity. Thus, RR CTL have the potential to kill infected macrophages but, if they cannot release granulysin, would likely release viable bacilli, resulting in disseminating the intracellular bacteria rather than controlling the infection. In human vaccine trials in tuberculosis there is a critical need for correlates and biomarkers of protection. The data here suggest that measurement of IFN-γ production by CD8^+^ T cells, most often used in vaccine trial recipients, will simply not serve as a useful measure of amCTL, which would need to be directly estimated by measurement of granulysin or its specific surface marker correlates ^22^.

Surprisingly, our analysis indicated that fibroblasts, keratinocytes and endothelial cells, cells not typically considered to contribute to the antimicrobial response, indeed expressed antimicrobial genes in RR granulomas. There were two distinct fibroblast subpopulations in RR: *CXCL2^+^* fibroblasts, described as inflammatory ^32^ and *SFRP2^+^* fibroblasts, involved in the deposition of extracellular matrix proteins ^31^. By spatial-seq, we identified *SFRP2^+^* fibroblasts in the papillary dermis of RR and T-lep lesions, just beneath the epidermis. In contrast, *CXCL2^+^* fibroblasts were located at the periphery of granuloma. Fibroblasts expressing vimentin are microanatomically located at the periphery of the tuberculosis granuloma ^42^, and are thought to sequester bacteria within the granuloma and prevent them from disseminating. We found that both fibroblast subpopulations express a number of chemokines that are also directly antimicrobial, hence they both represent fibroblast subpopulations with potential antimicrobial activity. Both fibroblast subpopulations contribute to the production of *CXCL9, CXCL10* and *CXCL11* that have been linked to trained immunity against mycobacterial infection in macrophages ^43^. *CXCL2^+^* fibroblasts also express *IL32,* which was shown to be involved in the vitamin D-dependent antimicrobial pathway and a marker of protective immunity in tuberculosis ^44^. We reason that the fibroblasts surrounding the granuloma form a dual function in host defense: forming a physical wall of extracellular matrix proteins and by the capacity to release antimicrobial proteins.

The epidermis is activated in RR, with hyperplasia of keratinocytes expressing MHC class II and IP-10, indicating activation by IFN-γ ^45,46^. Keratinocytes in RR expressed genes encoding antimicrobial peptides associated with antimycobacterial activity: *DEFB1* ^47^, *RNASE7* ^48^, *S100A8/S100A9* ^49–51^ and *TAC1* ^52^. Production of antimicrobial peptides from keratinocytes results in increased antimicrobial activity in the dermis, presumably by diffusion of the peptide across the dermal epidermal junction ^53^. The pseudotime mapped keratinocyte maturation, ending with a gene pattern indicating activation by both IL-1β and IFN-γ, with expression of *IL18,* which encodes for a protein that further upregulates IFN-γ in mycobacterial infection ^54^. Endothelial cells expressed a number of antimicrobial genes including *APP, CCL24, CXCL11, LEAP2, SNCA, TSLP, VIP, EC1, EC3, EC4, CCL21* and *NTS* ^36^.

Several features of the subtypes that were overrepresented in L-lep lesions may contribute to pathogenesis in leprosy. TREM2 macrophages, abundant in L-lep lesions, have been identified in several diseases characterized by altered lipid metabolism including atherosclerosis, Alzheimer’s disease, non-alcoholic steatohepatitis and obesity ^24–30^. The gene expression pattern in TREM2 macrophages suggests that these cells are programmed to transport and process lipids, most likely they are the foam cells or foamy macrophages that characterize both atherosclerosis and L-lep. One myeloid subtype, Type I IFN MΦ, and one T cell subtype, L-lep CTL, are both characterized by a type I IFN gene program. Previously, we found that IFN-β and its downstream targets were preferentially expressed in L-lep lesions and could inhibit IFN-γ induced antimicrobial activity in macrophages ^11^. We did not enrich for Langerhans cells from L-lep lesions; however they are less frequent in L-lep than in RR biopsy specimens ^3,55^ Tregs were present in both RR and L-lep lesions, and we were not able to identify CD8^+^ T suppressor cells in L-lep lesions, perhaps because of the low detection of *IL4.*

It has long been thought that the nature of the immune responses in infection, cancer and autoimmune diseases is dictated by the principal cells of the immune system, lymphocytes and myeloid cells. However, a compelling aspect of our data on leprosy is that not only these immune cells, but also other cell types such as fibroblasts and keratinocytes, cells that are not thought of as the traditional immune cells in the granulomatous immune response, have the capability of responding to *IL1B* and *IFNG* to produce antimicrobial molecules. Our data indicate that each major cell type, T cells, myeloid cells, fibroblasts and keratinocytes, expressed a unique set of antimicrobial genes. Furthermore, we have mapped the spatial location of these cell types, elucidating the granuloma architecture: a central core of differentiating macrophages with a mantle zone composed of dendritic cells and T cells, that express and *IL1B* and *IFNG,* respectively, key upstream regulators of the antimicrobial gene response and the macrophage psuedotime trajectory. Thus, the cells in the mantle zone have a key role in producing antimicrobial proteins to contain the bacteria but also via the release of cytokines in activating cells within and outside the granuloma to contribute to the antimicrobial response. The immune response in granulomas likely extends beyond its structural border to include keratinocytes and Langerhans cells in the epidermis, fibroblasts that accumulate near the epidermis and endothelial cells throughout the dermis. Our studies suggest a temporal and spatial model in which immunological activation of multiple cell types in and around the organized structure of the granuloma can lead to control of powerful intracellular pathogens. One could summarize our key findings simply by saying that it would appear to take a village to create effective antimicrobial granulomas.

## Acknowledgements

This work was supported in part by NIH grants AI-22553 and AR-40312 to R.L. M., NIH-P30 AR075043 (J.E.G), the Searle Scholars Program, the Beckman Young Investigator Program, a Sloan Fellowship in Chemistry, the NIH (5U24AI118672), and the Bill and Melinda Gates Foundation to A.K.S.

## Author Contributions

Conceptualization - F.M., T.K.H., B.B., A.S., B.R.B., M.P., R.L.M.; Methodology - F.M., T.K.H., R.M.B.T., P.R.A., B.J.A.S., O.P., A.C., T.D., M.H.W., M.P., R.L.M.; Investigation - F.M., T.K.H., R.M.B.T., P.R.A., B.J.A.S., T.D., M.T.O, E.N.S.; Formal Analysis - F.M., T.K.H., B.B., R.M.B.T., P.R.A., B.J.A.S., T.D., M.H.W.; Writing - F.M., T.K.H., R.M.B.T., P.R.A., B.J.A.S., M.H.W., A.O., M. L.I.-A., B.B., A.K.S., B.R.B., J.E.G., M.P., R.L.M.; Supervision - M.P., R.L.M.

## Declaration of Interests

A.K.S. reports compensation for consulting and/or SAB membership from Merck, Honeycomb Biotechnologies, Cellarity, Cogen Therapeutics, Orche Bio, and Dahlia Biosciences.

## Methods

### IRB Statement

Informed written consent was obtained from human subjects under a protocol approved by the institutional review boards of the Oswaldo Cruz Foundation, University of Southern California and University of California Los Angeles. This study was conducted according to the Declaration of Helsinki Principles.

### Processing of Human Skin

Skin biopsy specimens were obtained from patients with leprosy at University of Southern California and Oswald Cruz Foundation (Brazil). Patients were classified according to standard clinical and histologic criteria ^2^ (**Table 1**). Five patients with reversal reaction are designated here as RR1, RR2, RR3, RR4 and RR5. The other five are designated here as L-lep1, L-lep2, L-lep3, L-lep4 and L-lep5, of which four were classified as LL and one, L-lep2, as BL.

For each sample, a 4-mm punch biopsy was obtained following local anesthesia and was placed immediately into 10 mL of RPMI on ice. Initially, skin biopsies were incubated in 5mL of a 0.4% Dispase II solution (Roche Inc.) at 37°C for 1 hour with vigorous shaking. The dermis and epidermis were then carefully separated using forceps and transferred to separate tubes for additional processing. Epidermal samples were placed in 3mL of 0.25% Trypsin and 10U/mL DNAse for 30 minutes at 37°C. Trypsin was neutralized with 3mL of fetal calf serum (FCS), and the tissue was passed through a 70-micron nylon cell strainer which was washed with 5mL of RPMI. Epidermal cells were then pelleted at 300xg for 10 minutes and counted. Dermal samples were minced with a scalpel and incubated in a solution of 0.4% collagenase type II and 10 U/mL DNAse for 2 hours at 37°C with agitation. The cell suspension was passed through a 70-micron cell strainer and washed with 5mL of RPMI. Cells were pelleted at 300xg for 10 minutes, resuspended in 1mL of RPMI and counted. MACS enrichment for CD1a^+^ cells was performed for epidermis from three RR patients.

### Single cell library preparation

Once a single cell suspension was obtained, we utilized Seq-Well, a massively-parallel scRNA-Seq platform for clinical samples, to capture the transcriptome of single cells. Full methods on implementation of Seq-Well are available in Gierahn *et al.* ^20^ or on the Shalek Lab website (www.shaleklab.com). Briefly, 10-15,000 cells were loaded onto a functionalized-polydimethylsiloxane (PDMS) array preloaded with uniquely-barcoded mRNA capture beads (Chemgenes; MACOSKO-2011-10). After cells had settled into wells, the array was then sealed with a hydroxylated polycarbonate membrane with pore sizes of 10 nm, facilitating buffer exchange while confining biological molecules within each well. Following membrane-sealing, subsequent buffer exchange permits cell lysis, mRNA transcript hybridization to beads, and bead removal before proceeding with reverse transcription. The obtained bead-bound cDNA product then underwent Exonuclease I treatment (New England Biolabs; M0293M) to remove excess primer before proceeding with second strand synthesis.

### Sequencing and alignment

Libraries were sequenced on an Illumina Nova-Seq (Illumina, San Diego, CA) as 50bp paired end reads and were converted from bcl files to fastq files using bcl2fastq. We use Nextera N700 indices to identify individual samples. The alignment was performed using Drop-seq pipelines (version 1.12) previously described ^18^. Briefly, the raw reads were aligned to the concatenated human (hg38) and *M*. *leprae* genome using STAR ^56^. Each read was tagged with a 12bp barcode and 8bp unique molecular identifier (UMI). After alignment, the reads were grouped by the barcodes and deduplicated using the UMIs. The number of UMIs was then counted for each gene in each cell to generate the digital expression matrix (DEM).

### Removal of ambient RNA contamination

Ambient RNA contamination was removed using SoupX ^57^. Specifically, we examined the distribution of UMIs for each gene and selected the genes for which the distribution most closely approximated a uniform distribution. For each sample, we calculated an array-specific “soup” profile among barcodes below the UMI threshold. To calculate estimated per-cell contamination fractions, we manually selected genes observed to be bimodally expressed across cells, which suggest that these genes are predominantly expressed in a single cell type but are observed at low levels in other cell types for which endogenous expression would not be expected. For each array, we removed individual transcripts most likely to be contamination from each single cell based on the estimated contamination fraction. Specifically, individual transcripts were sequentially removed from each single cell transcriptome until the probability of subsequent transcripts being soup-derived was less than 0.5 to generate a background-corrected UMI matrix for each Seq-Well array.

### Cell clustering and cell type annotation

Digital expression matrices for human genes from all 10 samples were merged, and the R package Seurat ^21^ was used to cluster the cells in the merged matrix. Cells with less than 300 genes detected or more than 50% mitochondrial gene expression were first filtered out as low-quality cells. Genes detected in less than five cells were removed as low-abundance genes. The gene counts for each cell were divided by the total gene counts for the cell and multiplied by the scale factor 10,000, then natural-log transformation was applied to the counts. The FindVariableFeatures function was used to select 2,000 variable genes with default parameters. The ScaleData function was used to scale and center the counts in the dataset. Principal component analysis (PCA) was performed on the variable genes, and 13 PCs (based on the elbow point of variance explained by each PC) were used for cell clustering (resolution = 0.5) and Uniform Manifold Approximation and Projection (UMAP) dimensional reduction. The cluster markers were found using the FindAllMarkers function, and cell types were manually annotated based on the cluster markers. To generate the heatmap showing the cell type markers, the top 100 cells with the highest number of UMI detected were plotted for each cell type. The total number of *M. leprae* UMIs were calculated for each cell and plotted for each sample.

### Cell type sub-clustering

We performed sub-clustering on endothelial cells, fibroblasts, keratinocytes, myeloid cells and T cells. The same functions described above were used to obtain the sub-clusters. To choose the number of PCs, the rank of PCs based on the percentage of variance explained was plotted, and the elbow point was chosen as the number of PCs to use in cell clustering (resolution = 0.6) and UMAP dimension reduction. Clusters that were defined exclusively by mitochondrial gene expression, indicating low quality, were removed from further analysis. To generate the heatmap with marker genes for each sub-cluster, the top 100 sub-cluster marker genes with the highest average log fold change were plotted, and five representative genes were labelled.

### Interferon signature enrichment analysis

Supervised analyses were performed to identify Type I and Type II IFN regulated genes as described previously ^11,58–60^. Differentially expressed genes in TC1 (RR CTL) and TC2 (L-lep CTL) were identified using a Wilcoxon rank sum test with adjusted p value cutoff at 0.05. A list of genes specifically induced by only IFN-α/β or IFN-γ was derived from the gene expression profile data of IFN-treated human PBMC ^61^. 148 IFN-α/β specific genes and 33 IFN-γ specific genes were identified, which were overlapped with TC1 and TC2 specific genes to determine the differential expression of IFN-regulated genes. Hypergeometric test was used to determine the enrichment level; a p value smaller than 0.05 was considered to be significantly enriched.

### Pseudotime trajectory construction on macrophage and keratinocyte subtypes

Pseudo-time trajectories for macrophage and keratinocyte sub-clusters were constructed using the R package Monocle ^35^. The raw counts for cells in the intended sub-clusters were extracted and normalized by the estimateSizeFactors and estimateDispersions functions with the default parameters. Genes with average expression larger than 0.5 and detected in more than 10 cells were retained for further analysis. Variable genes were determined by the differentialGeneTest function with a model against the sub-cluster identities. The top 500 variable genes with the lowest adjusted p value were used to order the cells. The orders were determined by the orderCells function, and the trajectory was constructed by the reduceDimension function with default parameters. Differentially expression analysis was carried out using the differentialGeneTest function with a model against the pseudotime, and genes with an adjusted p value smaller than 0.05 were clustered into 6 patterns and plotted in the heatmap. Ingenuity Pathway Analysis was used to determine the upstream regulators for the genes in each expression pattern. A module score was calculated for each upstream regulator on gene targets from all six patterns. The module scores were calculated using the Seurat function AddModuleScore with default parameters, which measure the average expression levels of a set of genes, subtracted by the average expression of randomly selected control genes. Pearson correlation was then performed between the upstream regulator module scores and the pseudotimes.

### Antimicrobial gene analysis

A list of 1,404 genes were curated by searching for genes with “antimicrobial” as a keyword in GeneCards (https://www.genecards.org/). To study the difference of antimicrobial response in L-lep and RR, the cell types were split into L-lep and RR groups. To measure the relative abundance of anti-microbial genes (antimicrobial genes), the total expression of each antimicrobial gene was calculated for each L-lep and RR cell type. The antimicrobial gene expression for the L-lep cell types was normalized by the total number of L-lep cells, and the antimicrobial gene expression for the RR cell types was normalized by the total number of RR cells. The z scores were calculated across all L-lep and RR cell types for each antimicrobial gene. A cutoff of z score > 3 was applied to obtain the specific antimicrobial genes for each cell type. A list of 1,124 antimicrobial genes was obtained as specific to at least one RR cell type. Ingenuity Pathway Analysis was used to perform enrichment analysis and determine the upstream regulators of the 1,124 antimicrobial genes. To generate the circos plots, a list of direct antimicrobial genes was obtained from The Antimicrobial Peptide Database ^36^, and those regulated by *IL1B* or *IFNG* were included.

### Cell type composition analysis

To calculate the sample composition based on cell type, the number of cells for each cell type from each sample were counted. The counts were then divided by the total number of cells for each sample and scaled to 100 percent for each cell type. The same procedures were applied to calculate the sample composition for each subtype in endothelial cells, fibroblasts, keratinocytes, myeloid cells and T cells. The cell type (including the subtype) with more than 70% L-lep (or RR) composition was named L-lep (or RR) specific.

### Spatial sequencing library preparation

Skin samples were frozen in OCT medium and stored at −80°C until sectioning. Optimization of tissue permeabilization was performed on 20-μm-thick sections using Visium Spatial Tissue Optimization Reagents Kit (10X Genomics, Pleasanton, CA, USA), which established an optimal permeabilization time to be 6 minutes. Samples were mounted onto a Gene Expression slide (10X Genomics), fixed in ice-cold methanol, stained with hematoxylin and eosin, and scanned under a microscope (Keyence, Itasca, IL, USA). Tissue permeabilization was performed to release the poly-A mRNA for capture by the poly(dT) primers that are precoated on the slide and include an Illumina TruSeq Read, spatial barcode, and unique molecular identifier (UMI). Visium Spatial Gene Expression Reagent Kit (10X Genomics) was used for reverse transcription to produce spatially barcoded full-length cDNA and for second strand synthesis followed by denaturation to allow a transfer of the cDNA from the slide into a tube for amplification and library construction. Visium Spatial Single Cell 3’ Gene Expression libraries consisting of Illumina paired-end sequences flanked with P5/P7 were constructed after enzymatic fragmentation, size selection, end repair, A-tailing, adaptor ligation, and PCR. Dual Index Kit TT Set A (10X Genomics) was used to add unique i7 and i5 sample indexes and generate TruSeq Read 1 for sequencing the spatial barcode and UMI and TruSeq Read 2 for sequencing the cDNA insert, respectively.

### Spatial sequencing analysis

After sequencing, the reads were aligned to the human genome (hg38), and the expression matrix was extracted using the spaceranger pipeline. Seurat was then used to analyze the expression matrix. Specifically, the SCTransform function was used to scale the data and find variable genes with default parameters. PCA was applied for dimensional reduction. The FindTransferAnchors function was used to Find a set of anchors between the spatial-seq data and scRNA-seq data, which were then transferred from the scRNA-seq to the saptial-seq data using the TransferData function. Only the major cell types from the RR samples in the scRNA-seq data were used to annotate the spatial-seq data. The predicted cell type composition for each spot was then used to cluster the spots by the k-means algorithm. Six clusters were obtained for both the RR and T-lep spatial-seq sample. The clusters were annotated based the average cell type prediction score across all the spots in the cluster.

## Supplementary Figures

**Supplementary Fig. 1. Bacterial reads were mainly detected in the rRNA region in the *M. leprae* genome. Read alignment was shown in the interactive genome viewer for two patients, L-lep1 and L-lep2.**

**Supplementary Fig. 2: IFN-α/β and IFN-γ signature on CTL subtypes and co-expression of *IFNG, GZMB, PRF1* and *GNLY.***

a. Enrichment analysis on differentially expressed genes (adjusted p value < 0.05) between TC1 (RR CTL) and TC2 (L-lep CTL) using IFN-α/β and IFN-γ specific genes identified in human PBMC. Dotted lines indicate (left) no enrichment or (right) the hypergeometric test p value of 0.05 (log p value = 1.3).

b. UMAP plots showing co-expression of *IFNG, GMZB, PRF1* and *GNLY* in RR CTL and amCTL.

**Supplementary Fig. 3. Dendritic cell subpopulations and comparison of macrophage sub-clusters.**

a. UMAP plots showing *CD1C* and *LAMP3* expression in ML0. Only few co-expression events were observed, indicating distinct dendritic cell subpopulations.

b. Heatmap showing top differentially expressed genes between ML2 and ML4. ML3 expressed both ML2 and ML4 specific genes.

c. UMAP plots showing *TREM2* expression, *APOE* expression and TREM2 module score. The color scale represents the expression level of the genes or the level of the module score.

**Supplementary Fig. 4. Identification of keratinocyte subtypes.**

a. UMAP plot for 3,748 keratinocytes colored by subtypes.

b. UMAP plot colored by clinical forms.

c. Heatmap showing marker genes for each subtype. The representative genes are labelled.

d. Abundance composition across all samples for each keratinocyte subtype.

**Supplementary Fig. 5. Identification of fibroblast subtypes.**

a. UMAP plot for 1,010 fibroblasts colored by subtypes.

b. Abundance composition across all samples for each fibroblast subtype.

c. Heatmap showing marker genes for each subtype. The representative genes are labelled.

**Supplementary Fig. 6. Identification of endothelial cell subtypes.**

a. UMAP plot for 1,219 endothelial cells colored by subtypes.

b. Abundance composition across all samples for each endothelial cell subtype.

c. Dot plot showing 10 marker genes for each subtype. The color scale represents the scaled expression of the gene. The size of the dot represents percentage of cells expressing the gene.

**Supplementary Fig. 7. Pseudotime construction in macrophages and keratinocytes.**

a. Pseudo-temporal trajectory colored by pseudotime (top) and by sub-cluster identity (bottom) for macrophage sub-cluster 2, 3 and 4.

b. Heatmap showing six expression patterns along the macrophage pseudotime trajectory as depicted on the x axis. Representative genes regulated by *IL1B* and *IFNG* are labelled.

c. Pseudo-temporal trajectory colored by pseudotime (top) and by sub-cluster identity (bottom) for keratinocyte sub-cluster 1, 2 and 3.

d. Heatmap showing six expression patterns along the keratinocyte pseudotime trajectory as depicted on the x axis. Representative genes regulated by *IL1B* and *IFNG* are labelled.

**Supplementary Fig. 8. Representative antimicrobial genes expressed by myeloid cells, fibroblasts and keratinocytes.**

The height of the bar represents the z score of the gene in each cell type from RR lesions. The dots represent the gene’s expression level in individual cells.

**Supplementary Fig. 9. Representative antimicrobial genes expressed by Th17 cells, RR CTL and amCTL.**

The height of the bar represents the z score of the gene in each cell type from RR lesions. The dots represent the gene’s expression level in individual cells.

**Supplementary Fig. 10. Antimicrobial gene expression in distinct RR subtypes.**

Heat map showing z scores of antimicrobial genes in RR cell types. The red boxes indicate distinct sets of antimicrobial genes highly expressed in endothelial cells, fibroblasts, keratinocytes, myeloid cells and T cells.

**Supplementary Fig. 11. Cell type composition and clustering of the RR spatial-seq sample.**

a. Scatter pie plot showing the cell type composition of the RR spatial-seq sample. Each spot is represented as a pie chart showing the relative proportion of the cell types.

b. Heatmap showing the average cell type prediction score for each cluster.

**Supplementary Fig. 12. Spatial sequencing for the T-lep sample.**

a. H & E staining of the T-lep biopsy used for spatial sequencing.

b. Scatter pie plot showing the cell type composition of the RR spatial-seq sample. Each spot is represented as a pie chart showing the relative proportion of the cell types.

c. Spatial plot for 1,154 spots colored by clusters, the coordinates of the spot correspond to the location in the tissue.

d. Heatmap showing the average cell type prediction score for each cluster.

e. Spatial plots showing expression level of four antimicrobial genes. The colors correspond to the cluster colors in B. *CYBB* was highly expressed in myeloid cells, *CCL5* in T cells, *MMP2* in fibroblasts and *KLK5* in keratinocytes.

**Supplementary Fig. 13. Subtypes location resolved by spatial seq.**

a. Subtype prediction scores for the RR spatial-seq sample. The spots in the corresponding cluster was used to plot the subtype scores. For example, the FB0 prediction score was plotted in the spots from cluster 5, which was annotated as fibroblasts.

b. Subtype prediction scores for the T-lep spatial-seq sample. The spots in the corresponding cluster was used to plot the subtype scores.

## Supplementary Tables

**Supplementary Table 1. Clinical forms of each patient with leprosy.**

**Supplementary Table 2. Marker genes for each cell type from single cell RNA sequencing, and marker genes for the subtypes including T cells, myeloid cells, keratinocytes, fibroblasts and endothelial cells.**

**Supplementary Table 3. Type I and Type II Interferon signature genes derived from activated human PBMC.**

**Supplementary Table 4. Genes in six patterns along macrophage and keratinocyte pseudotime.**

**Supplementary Table 5. Antimicrobial genes curated from GeneCards and direct antimicrobial genes obtained from The Antimicrobial Peptide Database.**

**Supplementary Table 6. Validation of antimicrobial genes in leprosy and other mycobacterial infections as previously reported.**

**Supplementary Table 7. Marker genes for each cluster from spatial sequencing for the RR and T-lep sample.**

