## Supplementary Figures for "Single Cell and Spatial Transcriptomics Defines the Cellular Architecture of the Antimicrobial Response Network in Human Leprosy Granulomas"

### Supplementary Figure 1

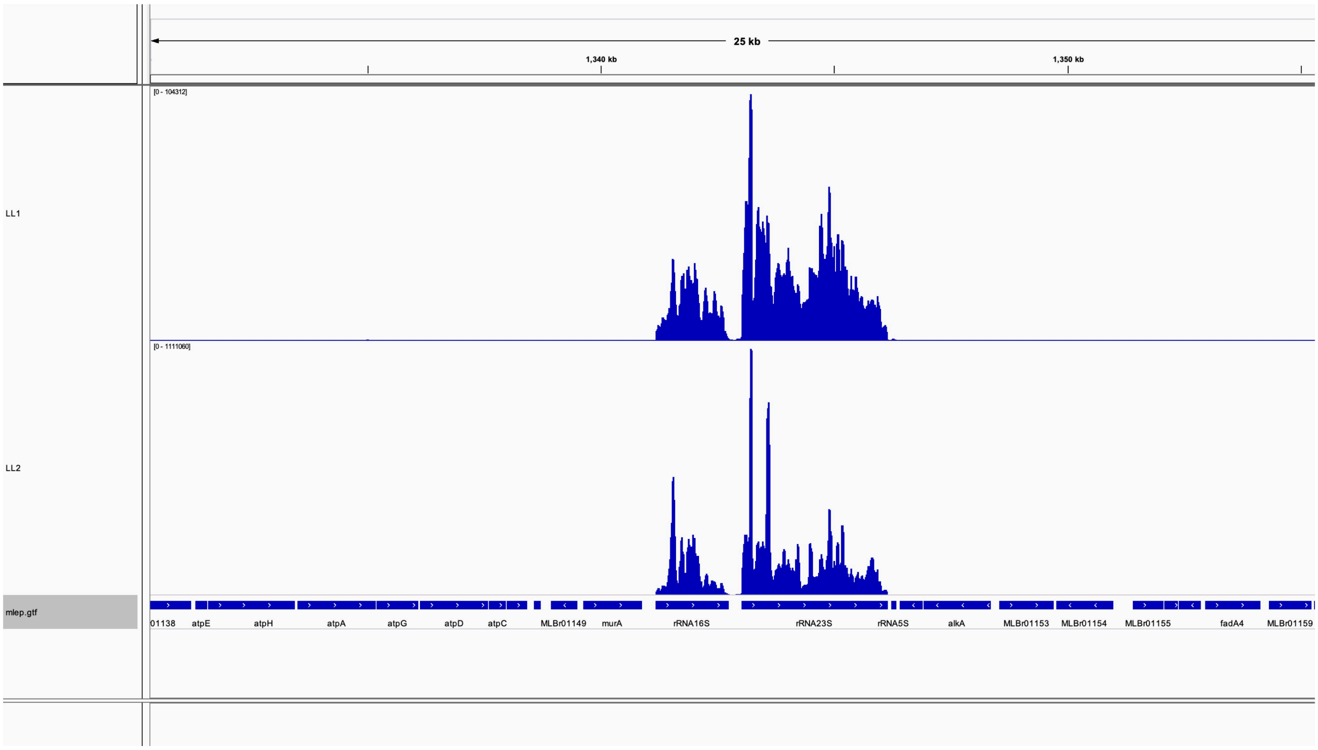

**Supplementary Figure 2**

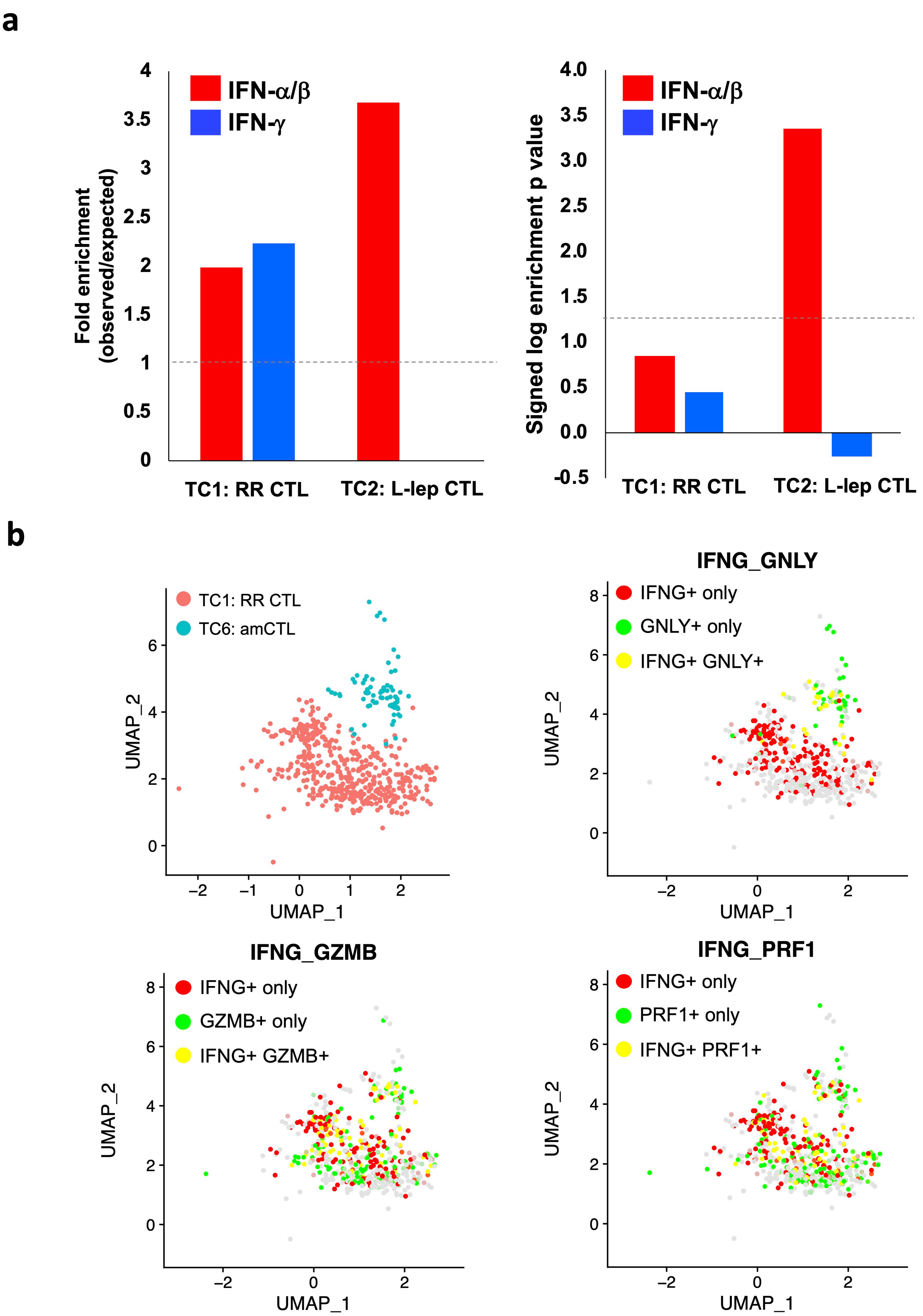

### Supplementary Figure 3

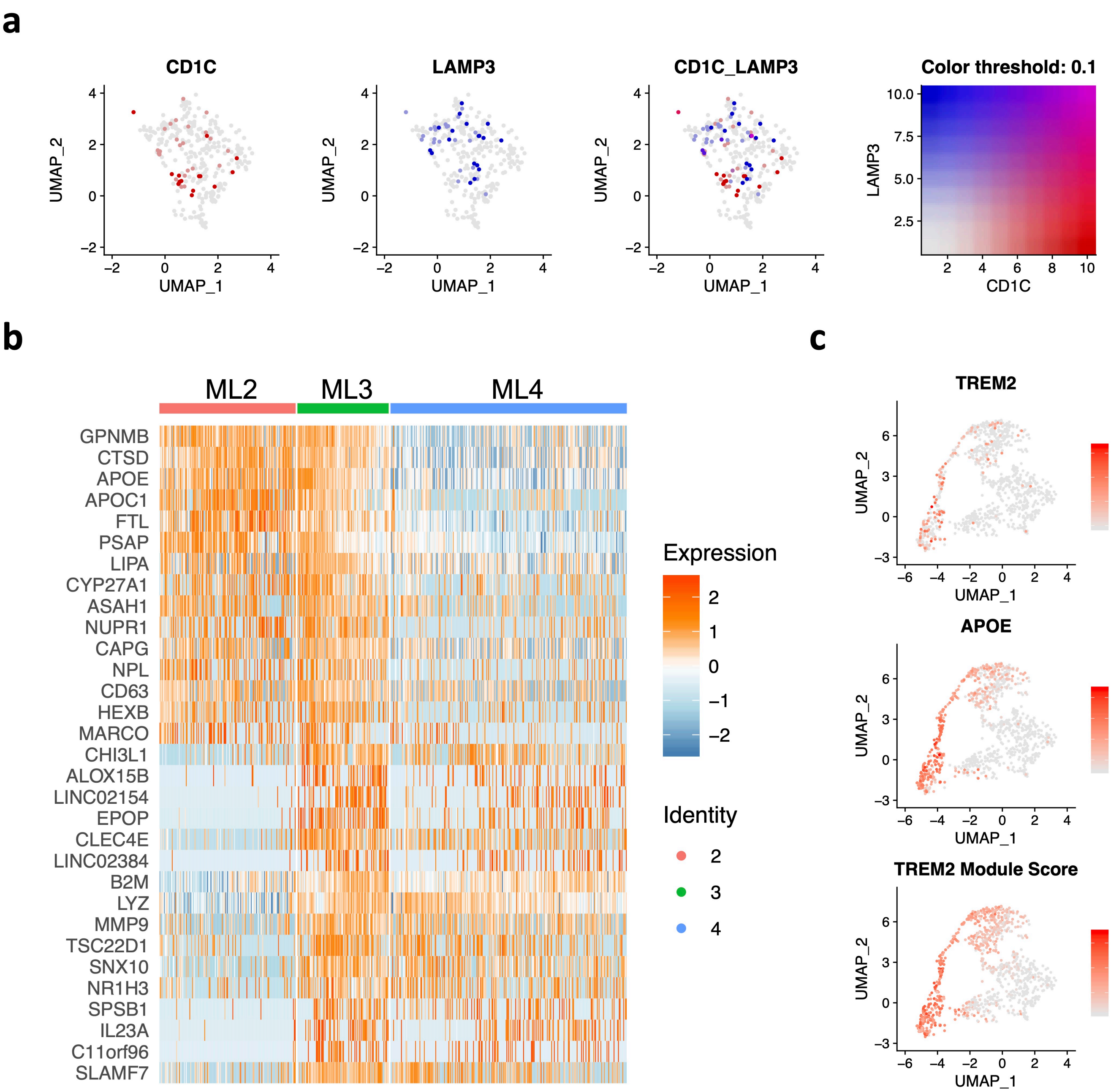

#### Supplementary Figure 4

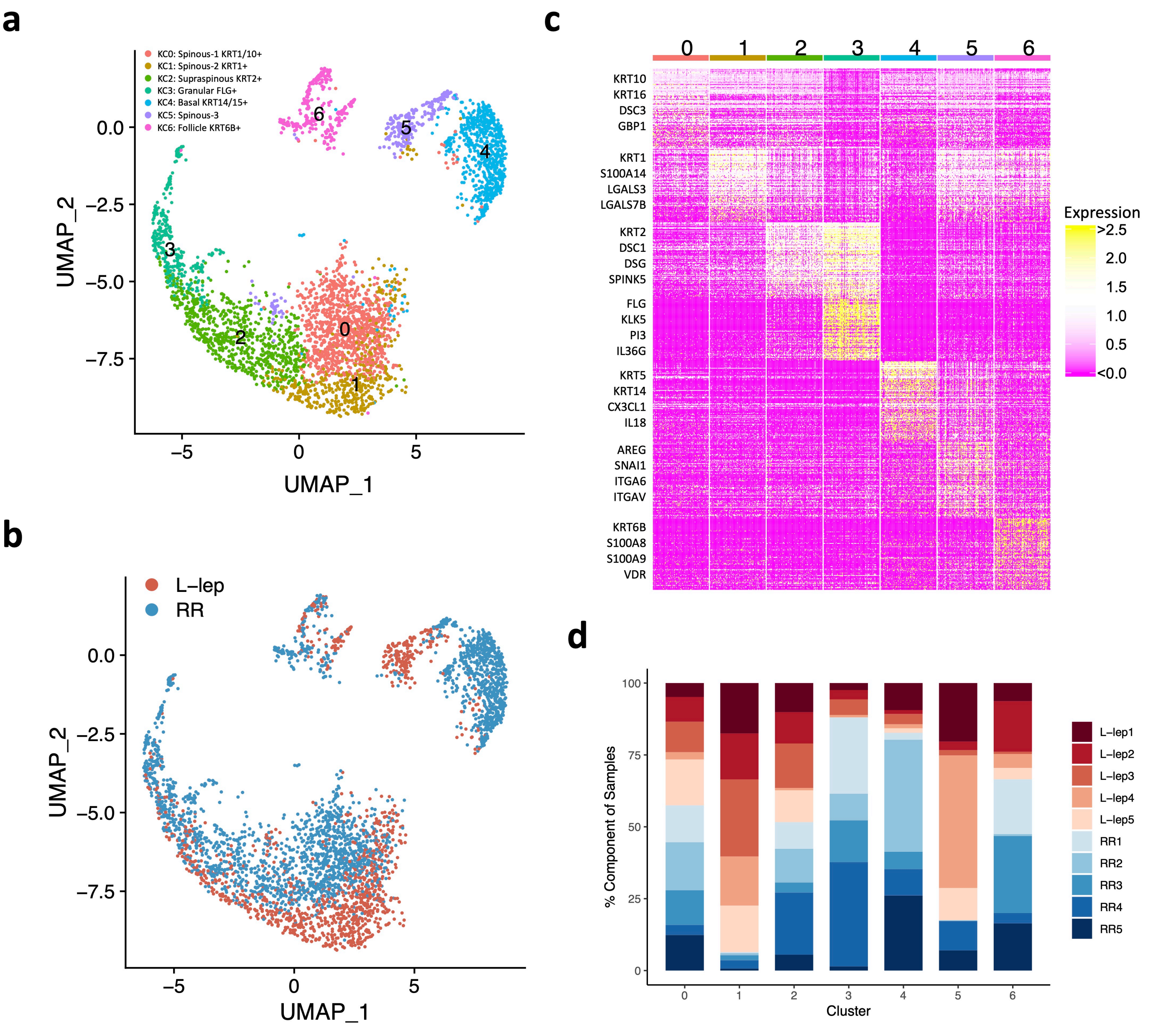

#### Supplementary Figure 5

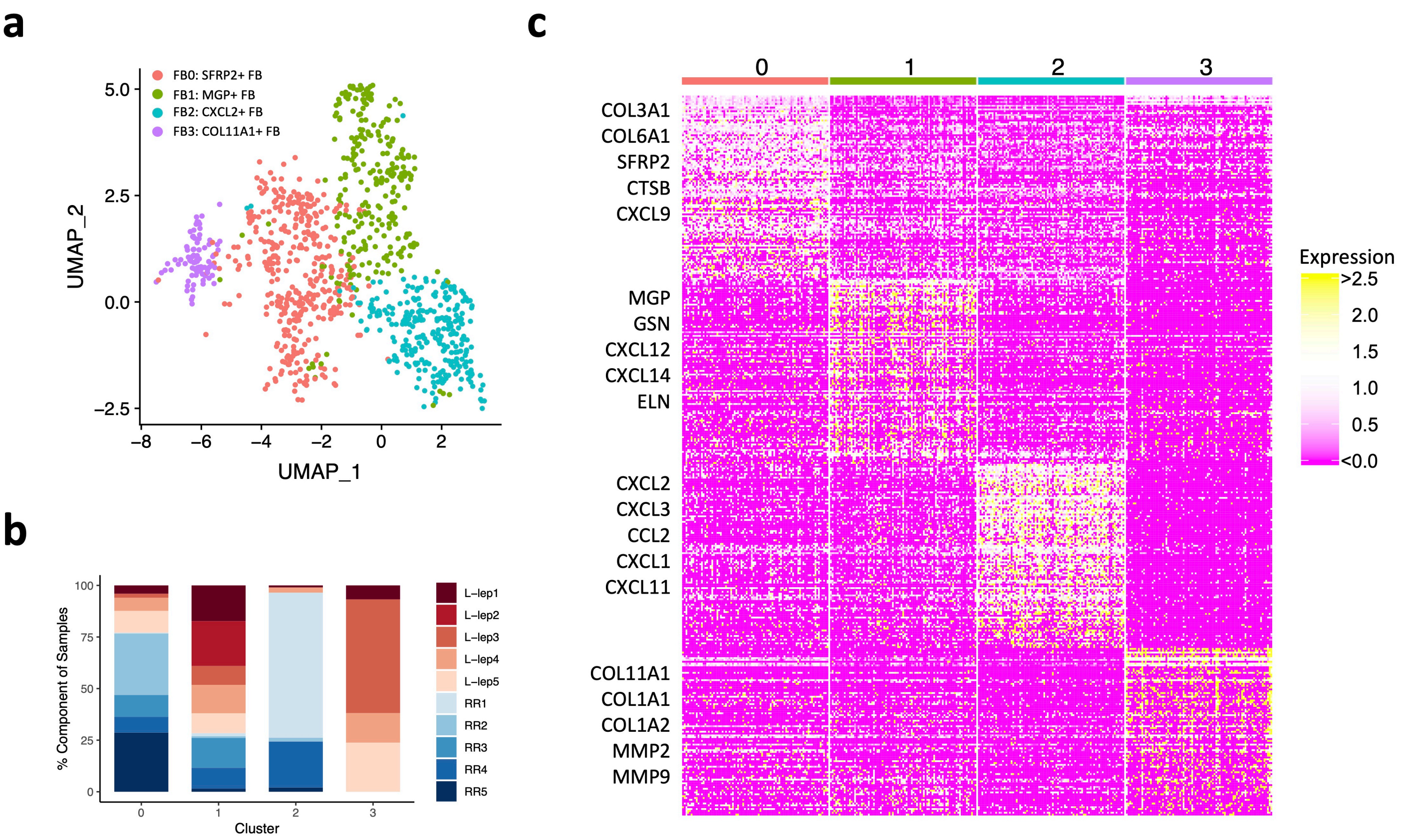

### Supplementary Figure 6

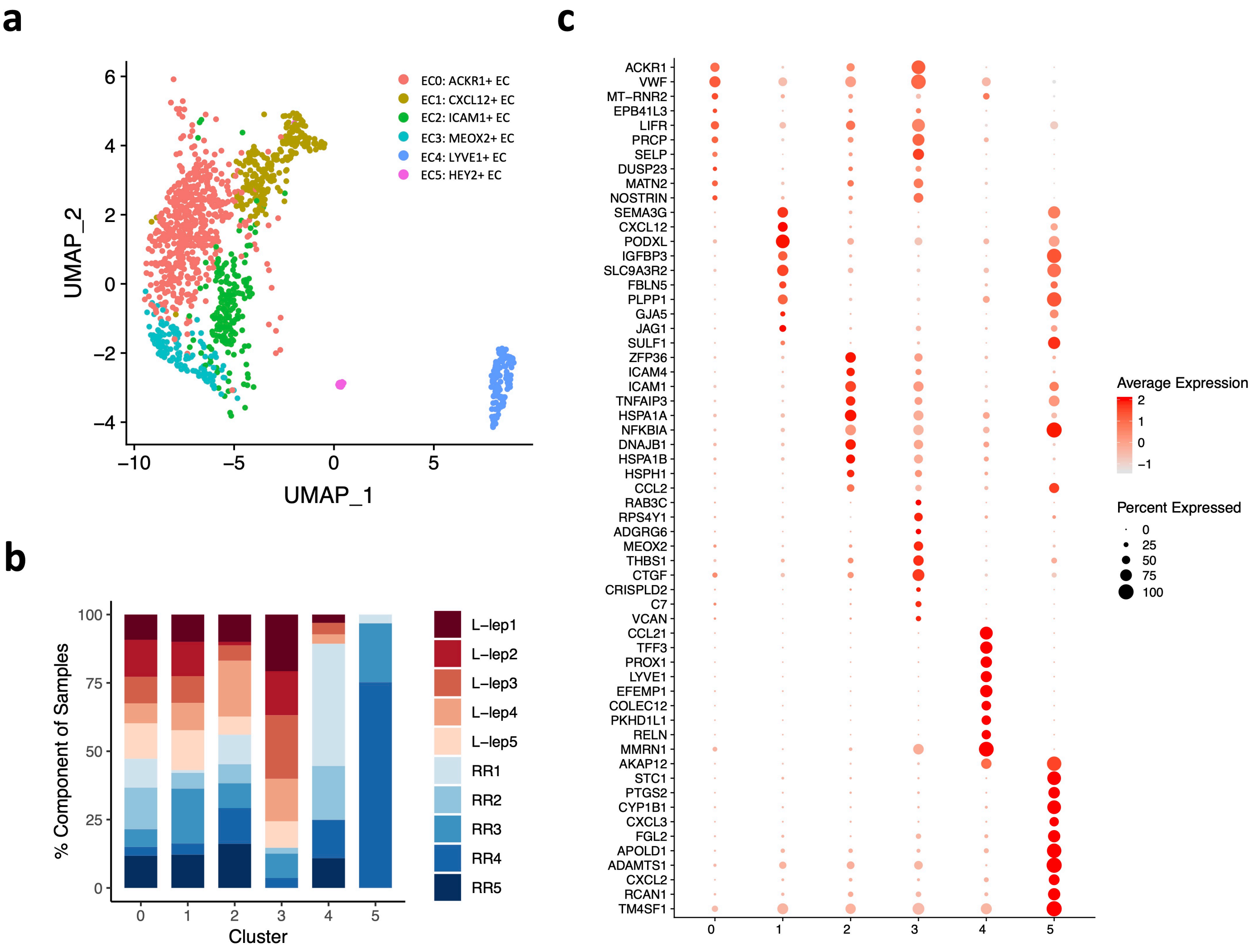

### Supplementary Figure 7

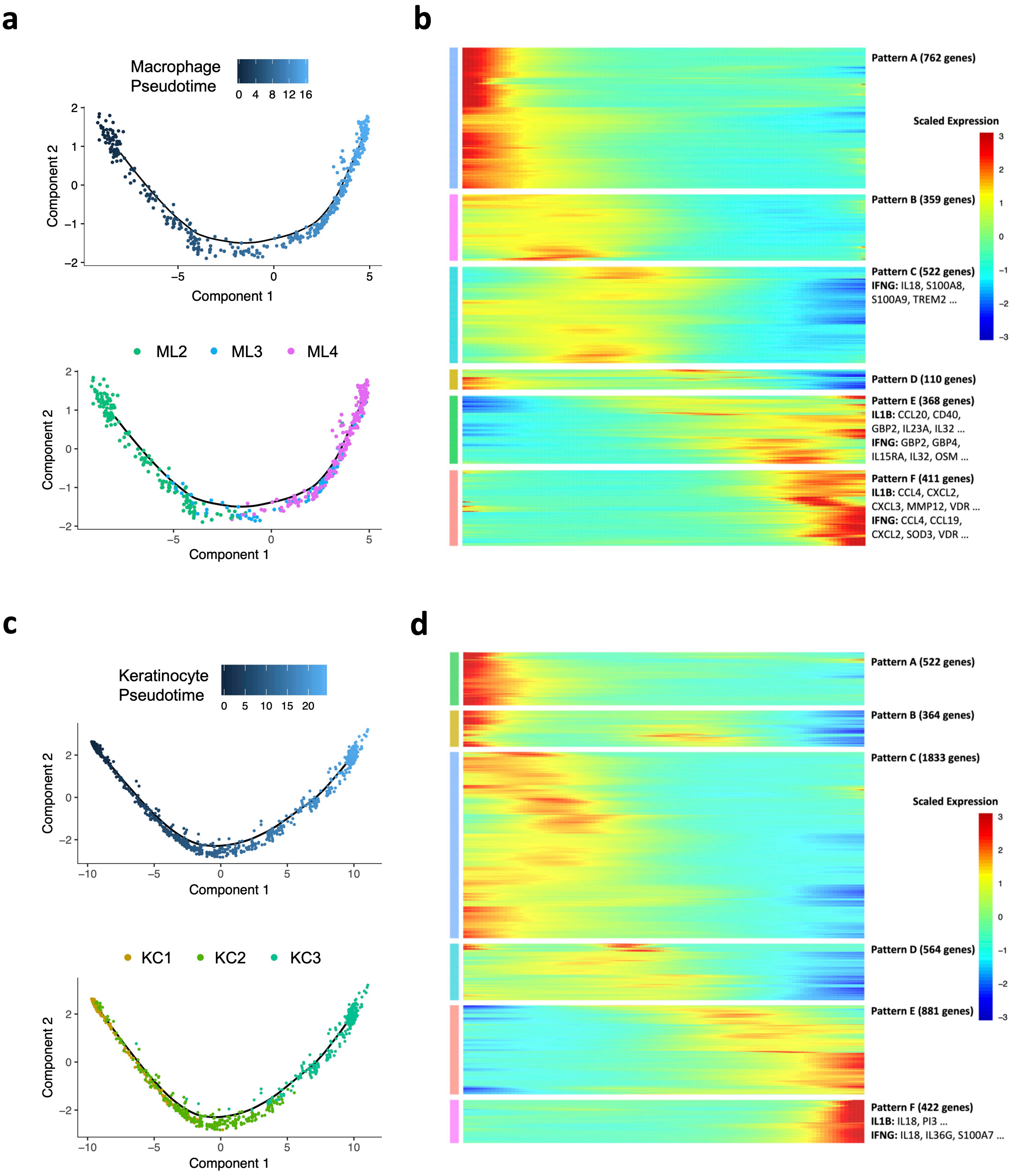

### Supplementary Figure 8

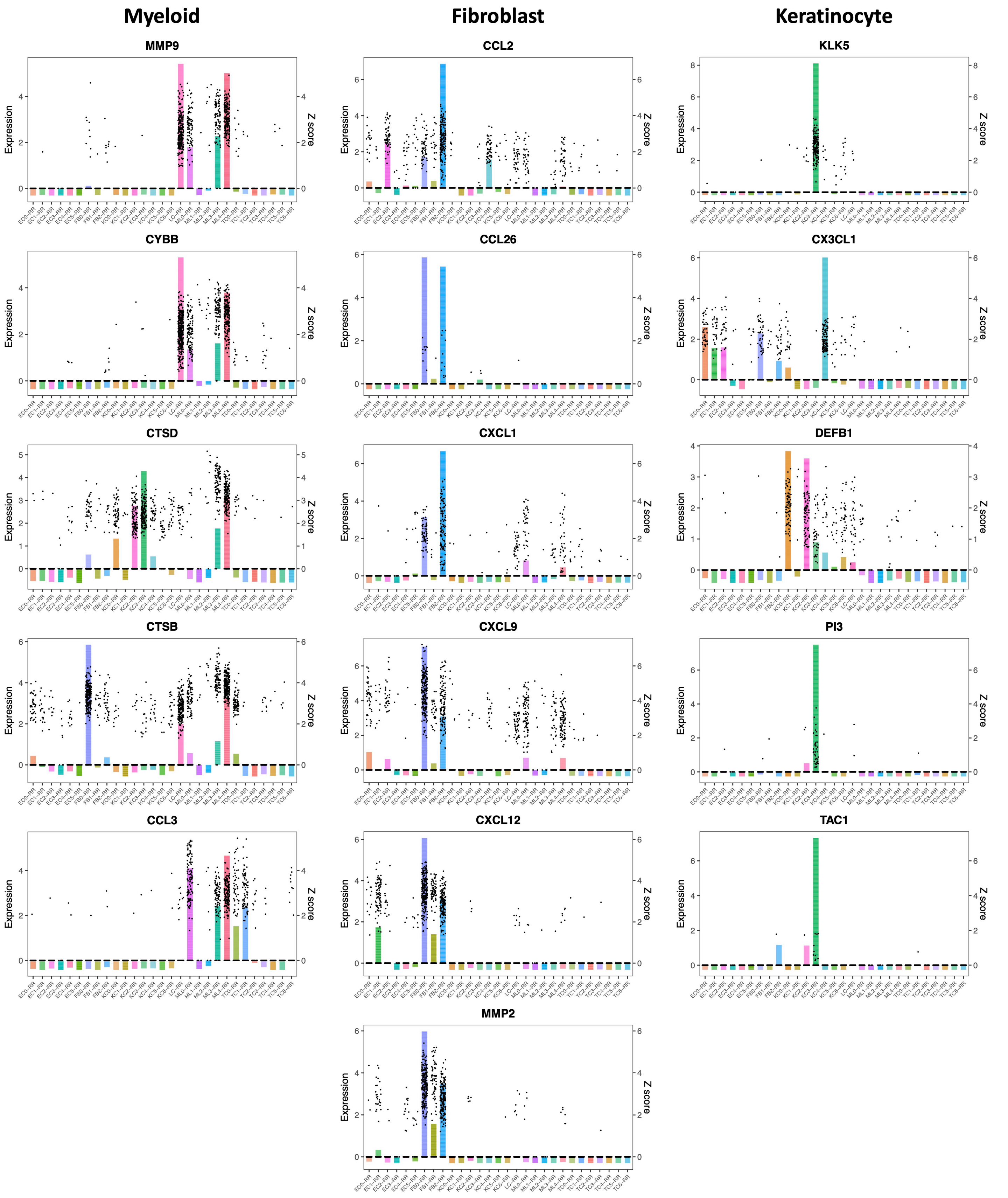

### Supplementary Figure 9

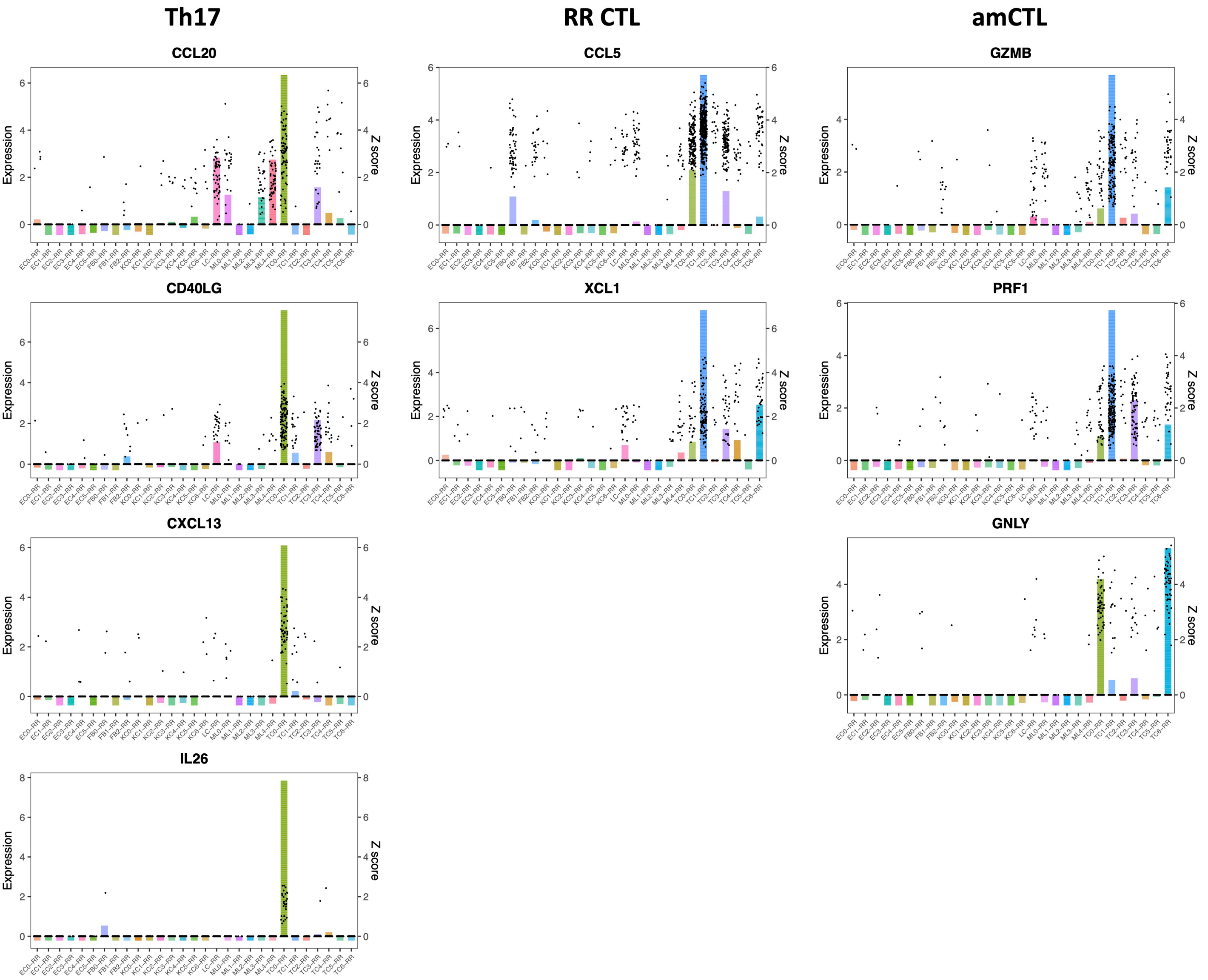

#### Supplementary Figure 10

**a**

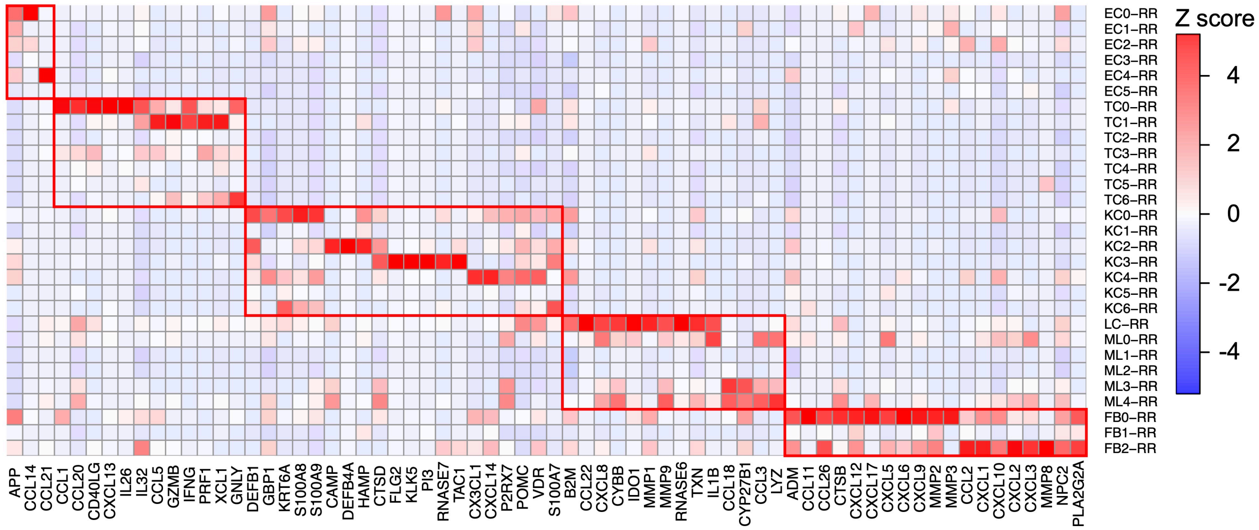

Supplementary Figure 11

**a**

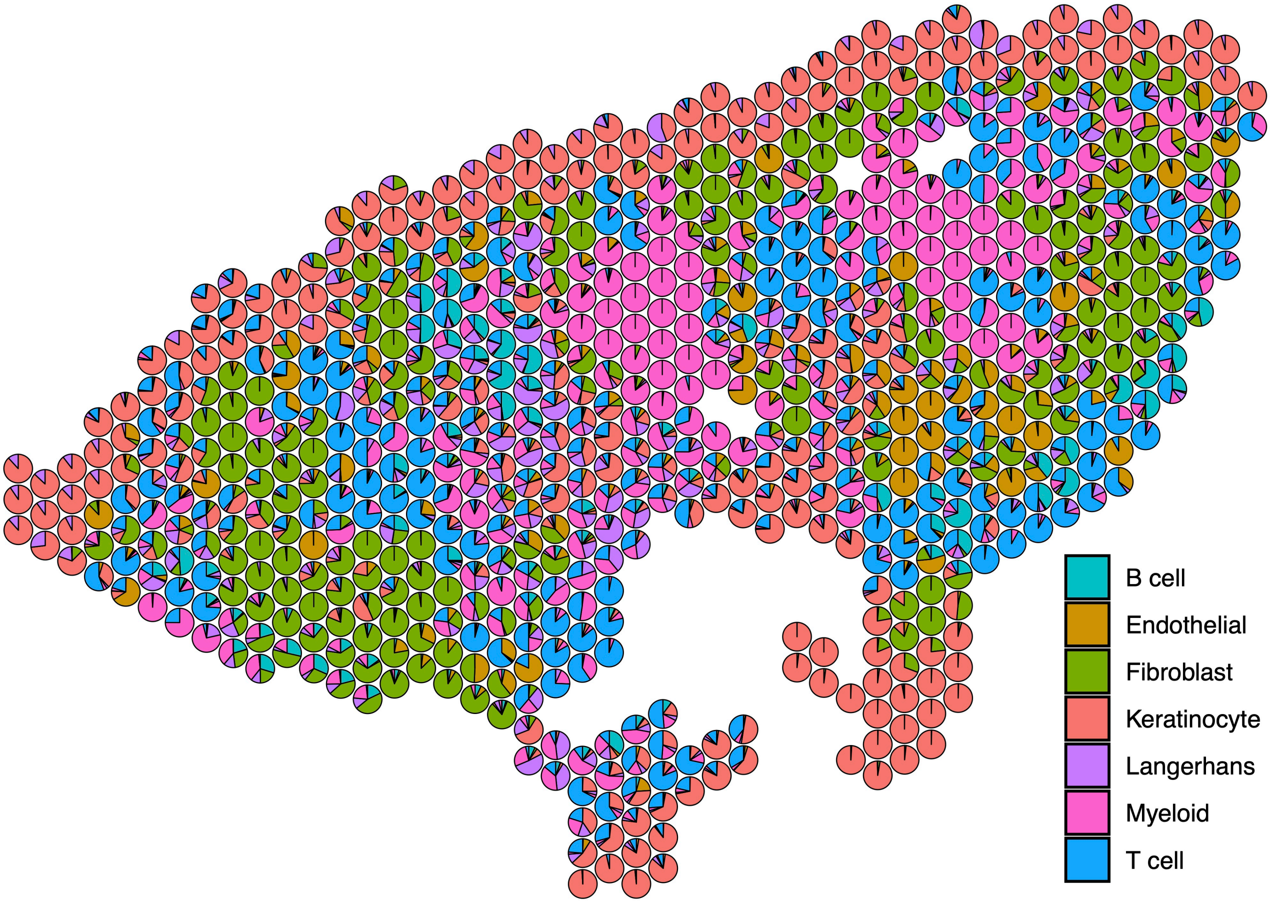


**b**

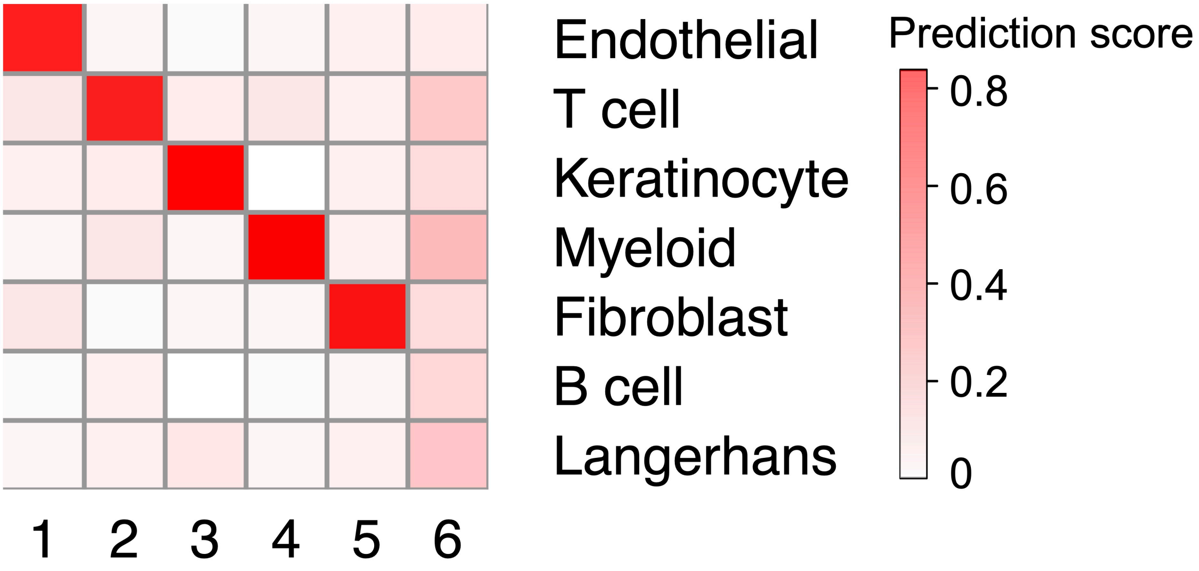

**Supplementary Figure 12**

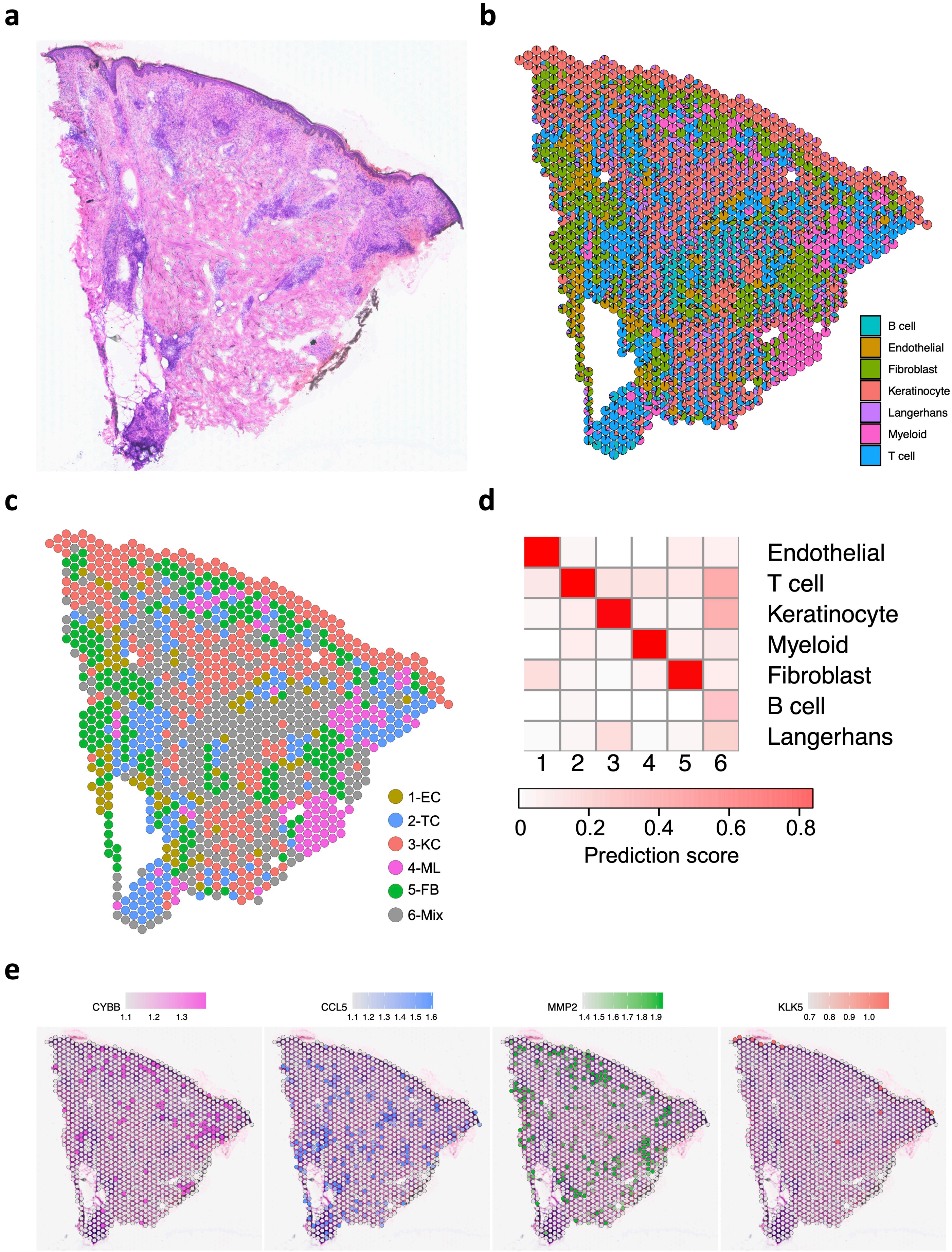

### Supplementary Figure 13

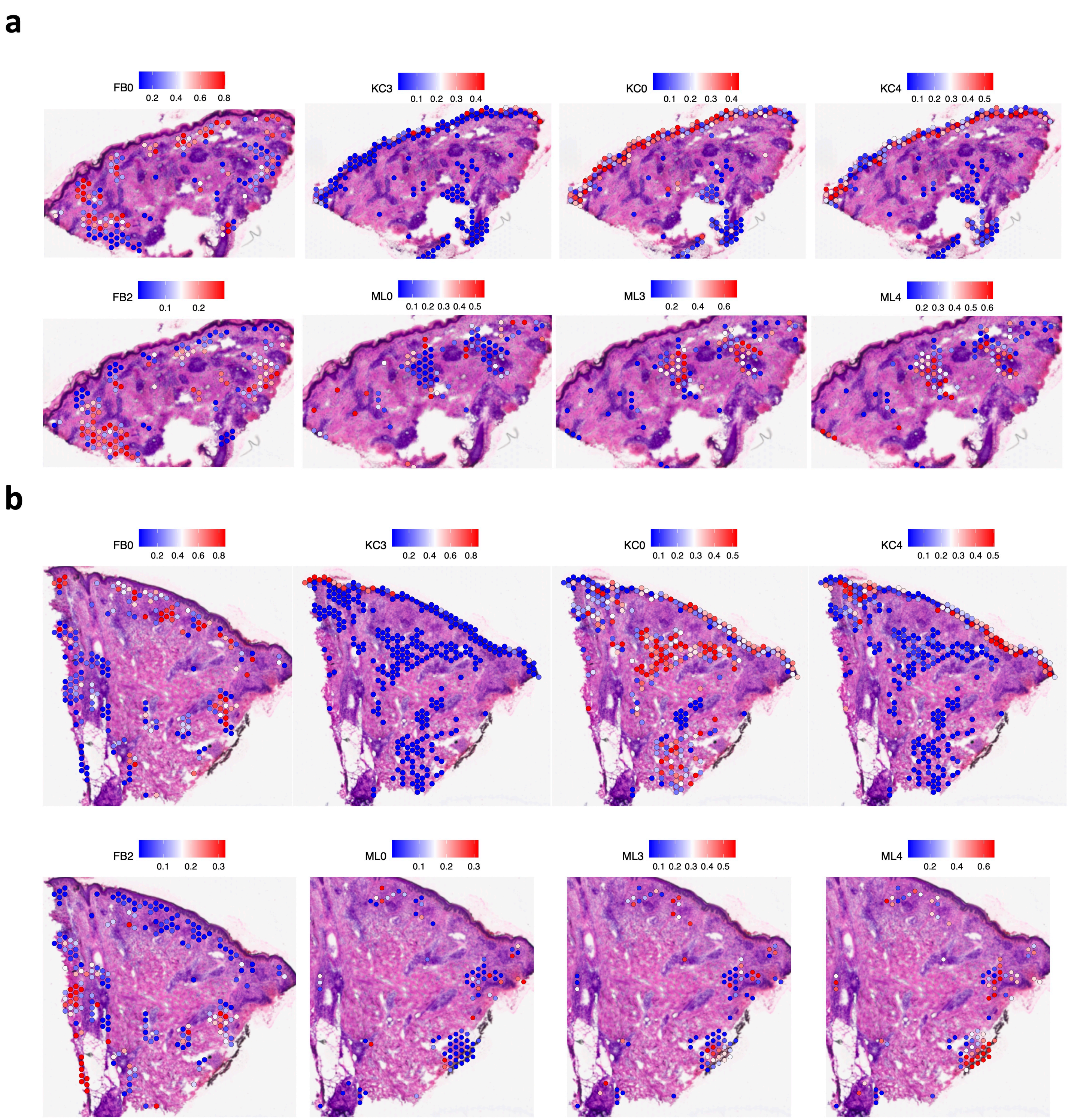
